# CD169-mediated restrictive SARS-CoV-2 infection of macrophages induces pro-inflammatory responses

**DOI:** 10.1101/2022.03.29.486190

**Authors:** Sallieu Jalloh, Judith Olejnik, Jacob Berrigan, Annuurun Nisa, Ellen L Suder, Hisashi Akiyama, Maohua Lei, Sanjay Tyagi, Yuri Bushkin, Elke Mühlberger, Suryaram Gummuluru

## Abstract

Exacerbated and persistent innate immune response marked by pro-inflammatory cytokine expression is thought to be a major driver of chronic COVID-19 pathology. Although macrophages are not the primary target cells of SARS-CoV-2 infection in humans, viral RNA and antigens in activated monocytes and macrophages have been detected in post-mortem samples, and dysfunctional monocytes and macrophages have been hypothesized to contribute to a protracted hyper-inflammatory state in COVID-19 patients. In this study, we demonstrate that CD169, a myeloid cell specific I-type lectin, facilitated ACE2-independent SARS-CoV-2 fusion and entry in macrophages. CD169- mediated SARS-CoV-2 entry in macrophages resulted in expression of viral genomic and sub-genomic (sg) RNAs with minimal viral protein expression and no infectious viral particle release, suggesting a post-entry restriction of the SARS-CoV-2 replication cycle. Intriguingly this post-entry replication block was alleviated by exogenous ACE2 expression in macrophages. Restricted expression of viral gRNA and sgRNA in CD169^+^ macrophages elicited a pro-inflammatory cytokine expression (TNFα, IL-6 and IL-1β) in a RIG-I, MDA-5 and MAVS-dependent manner, which was suppressed by remdesivir pre- treatment. These findings suggest that *de novo* expression of SARS-CoV-2 RNA in macrophages contributes to the pro-inflammatory cytokine signature and that blocking CD169-mediated ACE2 independent infection and subsequent activation of macrophages by viral RNA might alleviate COVID-19-associated hyperinflammatory response.

**Author Summary:** Over-exuberant production of pro-inflammatory cytokine expression by macrophages has been hypothesized to contribute to severity of COVID-19 disease. Molecular mechanisms that contribute to macrophage-intrinsic immune activation during SARS-CoV-2 infection are not fully understood. Here we show that CD169, a macrophage-specific sialic-acid binding lectin, facilitates abortive SARS-CoV-2 infection of macrophages that results in innate immune sensing of viral replication intermediates and production of proinflammatory responses. We identify an ACE2-independent, CD169- mediated endosomal viral entry mechanism that results in cytoplasmic delivery of viral capsids and initiation of virus replication, but absence of infectious viral production. Restricted viral replication in CD169^+^ macrophages and detection of viral genomic and sub-genomic RNAs by cytoplasmic RIG-I-like receptor family members, RIG-I and MDA5, and initiation of downstream signaling via the adaptor protein MAVS, was required for innate immune activation. These studies uncover mechanisms important for initiation of innate immune sensing of SARS-CoV-2 infection in macrophages, persistent activation of which might contribute to severe COVID-19 pathophysiology.

## Introduction

Severe acute respiratory syndrome coronavirus 2 (SARS-CoV-2) is the causative agent of the COVID-19 pandemic, which has claimed nearly 6 million deaths worldwide (https://coronavirus.jhu.edu/). Severe COVID-19 cases have been associated with aberrant bronchioalveolar immune cell activation and persistently high levels of proinflammatory cytokines, including IL-6, TNFα, and IL-1β (1, 2). This protracted immune hyperactivation state marked by uncontrolled proinflammatory cytokine expression (3–6) is a potential driver of acute respiratory distress syndrome (ARDS) in severe COVID-19. Transcriptomic analysis of bronchioalveolar lavage fluid (BALF) samples from SARS-CoV-2 infected individuals revealed extensive lung infiltration by inflammatory monocytes and activated tissue-resident and BALF-associated macrophages with robust induction of interferon-stimulated gene (ISG) expression (7), suggestive of a myeloid cell-intrinsic cytokine signature contributing to ARDS and COVID-19 pathologies (8, 9). However, whether SARS-CoV-2 can establish productive infection in monocytes and macrophages has remained contentious (10–15), and importantly, the molecular mechanisms that contribute to myeloid cell-intrinsic hyperinflammatory phenotype have remained unclear (6, 16–19).

Studies on post-mortem tissues from patients, who succumbed to COVID-19, showed that a subset of tissue-resident alveolar macrophages are enriched in SARS-CoV-2 RNA (20, 21). Additionally, single-cell RNA-seq analysis revealed the presence of viral mRNAs in inflammatory myeloid cell populations in autopsied lung tissues (22, 23). Recent studies suggest that tissue-resident human macrophages are permissive to SARS-CoV-2 infection in humanized mice models, and that inhibition of viral genome replication or type-I interferon (IFN) signaling significantly attenuates chronic macrophage hyperactivation and disease progression (24). However, whether the presence of viral RNA in macrophages reflects phagocytosis of infected bystander cells or active virus replication in tissue-resident macrophages has yet to be defined. In contrast, CD14^+^ peripheral blood monocytes, monocyte-derived dendritic cells (MDDCs), or monocyte-derived macrophages (MDMs) were not-permissive to productive SARS-CoV-2 replication in vitro (10–12, 15). In permissive lung epithelial cells and those expressing the cognate entry receptor, angiotensin-converting enzyme 2 (ACE2), SARS-CoV-2 utilizes its spike (S) glycoprotein to interact with ACE2, which facilitates proteolytic cleavage, plasma or endosomal membrane fusion, and cytosolic import of viral genome (25–27). Depending on cell type, different host proteases such as furin, TMPRSS2, or cathepsins are required for S cleavage and entry of SARS-CoV-2 (26, 28, 29). While circulating monocytes and macrophages are not known to express ACE2 (30), these cells have been shown to express low levels of endogenous surface TMPRSS2 (31), and moderate levels of endosomal cathepsins (32) (33), although the relative expression of these cellular proteases in the context of SARS-CoV-2 infection and inflammation is not well understood.

Recent reports have highlighted capture of SARS-CoV-2 virus particles by myeloid cell-specific receptors, such as C-type lectins, in an ACE2-independent manner (16, 17, 19, 34, 35) though virus particle fusion or productive viral infection was not observed. We and others have previously shown that CD169/Siglec-1 facilitates viral infections of macrophages or dendritic cell (DC)-mediated trans infection of bystander cells (36–39). CD169 binds to sialylated viral glycoproteins or viral membrane-associated gangliosides, GM1 and GM3 (38–44). SARS-CoV-2 S protein is highly sialylated (45, 46), and a recent report demonstrated that DC-mediated SARS-CoV-2 trans infection of ACE2^+^ epithelial cells was facilitated by CD169 (19). CD169 is highly expressed by splenic red pulp and perifollicular macrophages, subcapsular sinus macrophages (47) and alveolar macrophages (48, 49). Besides constitutive expression on tissue-resident macrophages, CD169 expression can be upregulated on peripheral blood monocytes under inflammatory conditions, especially in response to type I interferons (IFNs) (50–52). Since type I IFNs are highly upregulated and CD169-expressing myeloid cells are elevated during SARS-CoV-2 infection (53, 54), we reasoned that SARS-CoV-2 S mediated interactions with CD169^+^ macrophages might play a crucial role in driving immunopathology of SARS-CoV- 2 infection.

In this study, we examined the role of CD169 in facilitating SARS-CoV-2 infection of ACE2-deficient human macrophages and its effect on inducing pro-inflammatory cytokine expression. Using two different human macrophage models, PMA-differentiated THP1 cells (THP1/PMA) and primary monocyte-derived macrophages (MDMs), we show that CD169 binds to SARS-CoV-2 S and mediates SARS-CoV-2 S-dependent viral entry into macrophages, leading to restricted cytosolic expression of viral genomic and sub- genomic (sg) RNA. Surprisingly, induced constitutive expression of ACE2 in macrophages (THP1/PMA and MDMs) restored permissiveness to robust virus replication and production of infectious progeny virions, suggesting that ACE2 expression overcomes a post-entry block to SARS-CoV-2 infection in macrophages. While CD169-mediated, ACE2-independent SARS-CoV-2 entry into macrophages resulted in negligible viral protein expression and absence of infectious virus production, restricted expression of viral negative strand RNA and sgRNAs induced pro-inflammatory cytokine expression via retinoic acid-inducible gene I (RIG-I) and melanoma differentiation-associated gene 5 (MDA-5) dependent sensing of viral replication intermediates. Importantly, expression of IL-6, TNFα and IL-1β was enhanced, whereas type I IFN responses were muted, suggesting a novel CD169-mediated, macrophage-intrinsic amplification of pro-inflammatory responses. These findings suggest that induction of pro-inflammatory responses in SARS-CoV-2-exposed macrophages requires initial viral RNA synthesis and that abortively infected macrophages might contribute to the hyperimmune phenotype and pathophysiology of COVID-19.

## Results

### SARS-CoV-2 Spike protein can mediate ACE2-independent entry into macrophages

To examine the role of macrophages in SARS-CoV-2 infection and COVID-19 pathogenesis, we differentiated primary MDMs from multiple donors by culturing CD14^+^ monocytes in the presence of human AB-serum and M-CSF for 6 days (55). Compared to a control HEK293T/ACE2 cell line retrovirally transduced to stably express human ACE2, ACE2 expression in primary human MDMs was under the detection limit (**Fig. 1A and B**). However, similar to HEK293T/ACE2 cells (**Fig. 1C**), MDMs from multiple donors were robustly infected with a SARS-CoV-2 S-pseudotyped lentivirus (**Fig. 1D**), suggesting that macrophages can support ACE2-independent S-pseudotyped virus entry. Furthermore, SARS-CoV-2 S-pseudotyped infections in MDMs were blocked by pre-treatment with a cathepsin inhibitor (E64D) but not a TMPRSS2 inhibitor (Camostat) (**Fig. 1D**), suggesting that SARS-CoV-2 S facilitates endosomal viral entry into ACE2-deficient MDMs. Preferential engagement of endosomal entry mechanism for SARS-CoV-2 S pseudotyped lentiviruses (LVs) in MDMs correlated with lack of active and cleaved form of serine protease TMPRSS2 expression in THP1/PMA and primary macrophages ((56), **Fig. S1A**), and robust cathepsin-L expression in THP1/PMA and MDMs (**Fig. S1B**). Strikingly, pre-treatment with neutralizing antibodies targeting the N-terminal domain (NTD) of SARS-CoV-2 S that do not compete with ACE2 binding by S (57), led to significant reduction in S-pseudotyped lentivirus infection of primary MDMs, suggesting that specific interaction between SARS-CoV-2 S and ACE2-independent entry factors are essential to mediate entry and endosomal fusion in macrophages (**Fig. S2**).

**Figure 1.**
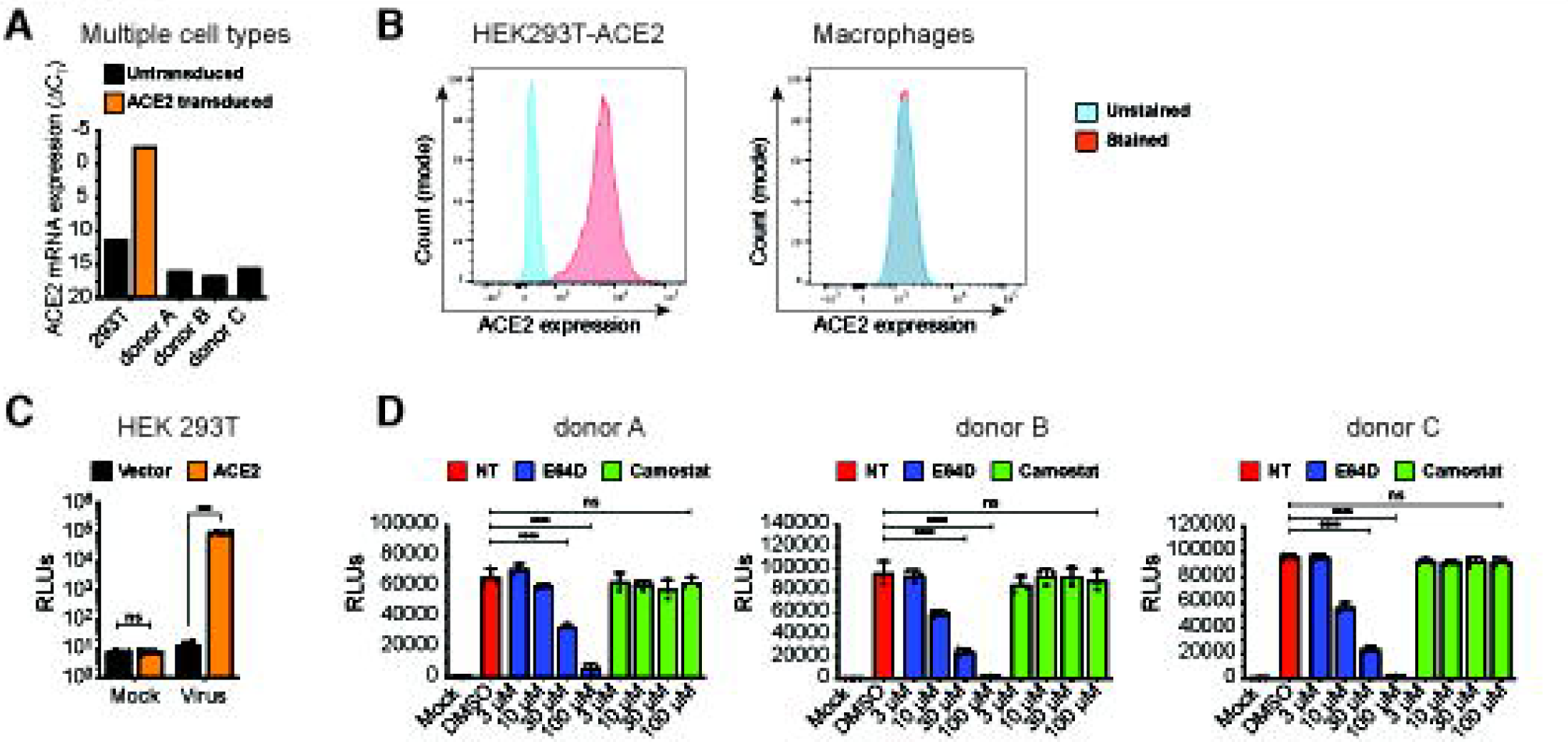
ACE2-independent SARS-CoV-2 entry in macrophages. (**A-B**) Representative ACE2 mRNA expression (**A**) and flow cytometry profiles showing ACE2 surface expression (**B**) of HEK293T cells stably expressing ACE2 and primary MDMs from multiple donors. (**C-D**) Parental (vector) and transduced (ACE2) HEK293T cells (**C**) and primary MDMs from 3 donors (**D**) were infected with S-pseudotyped lentivirus (20 ng based on p24^Gag^), in the absence or presence of cathepsin inhibitor (E64D) or TMPRSS2 inhibitor (Camostat), and infection was quantified by measuring luciferase activity at day 3 post-infection. Mock: no virus added, DMSO: no-treatment. The means ± SEM from at least 3 independent experiments are shown. *P*-values: paired t-test, two-tailed comparing to vector control (**C**), or one-way ANOVA followed by the Dunnett’s post-test comparing to DMSO (**D**). ***: *p* < 0.001, ****: *p* < 0.0001, ns: not significant.

### CD169 is a SARS-CoV-2 attachment and entry factor in macrophages

Expression of CD169, an ISG, is significantly upregulated in monocytes and alveolar macrophages isolated from COVID-19 patients, and its expression enhancement correlates with COVID-19 disease severity (7, 53). While co-expression of CD169 with ACE2 can enhance ACE2-mediated SARS-CoV-2 S entry in HEK293T cells (16), whether CD169 plays a role in SARS-CoV-2 infection of ACE2-deficient macrophages is unclear. To address this question, THP1 cells stably expressing wildtype (wt) CD169, mutant CD169/R116A which displays an attenuated ability to bind sialylated glycoconjugates (39, 58, 59), ACE2, or both wt CD169 and ACE2 (CD169/ACE2) (**Fig. S3A**) were incubated with recombinant SARS-CoV-2 S protein, and relative S binding was determined by flow cytometry. In contrast to parental THP1 monocytes, THP1 cells expressing wt CD169 (THP1/CD169) displayed robust S binding comparable to levels observed with THP1/ACE2 cells (**Fig. 2A**). There was a significant reduction in S binding to THP1/CD169-R116A cells, indicating that the sialylated S protein is recognized by CD169 (**Fig. 2A**). Co-expression of CD169 and ACE2 in THP1 cells further enhanced S binding compared to cells expressing only ACE2 or CD169, suggestive of cooperative binding of CD169 and ACE2 to SARS-CoV-2 S.

**Figure 2.**
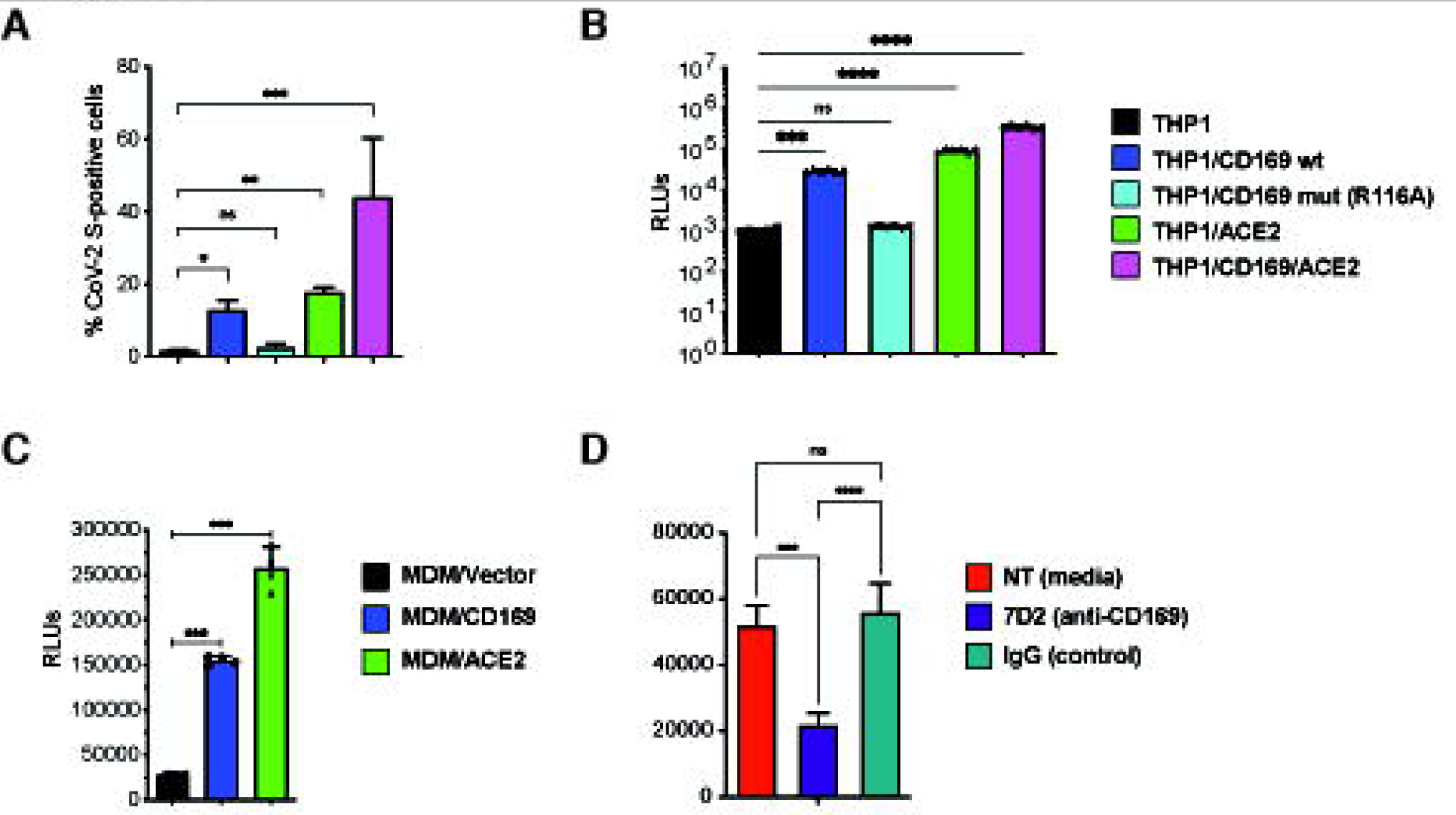
CD169 is a SARS-CoV-2 attachment and entry factor in macrophages. (**A**) Binding of SARS-CoV-2 S protein (Wuhan isolate) to THP1 monocytes expressing wt CD169, mutant CD169 (R116A), ACE2, or both wt CD169 and ACE2 (CD169/ACE2). (**B**) THP1/PMA macrophages were infected with S-pseudotyped lentivirus (20 ng p24^Gag^), and infection was quantified by measuring luciferase activity at 3 dpi. Relative light units (RLUs) from each cell line were normalized to no virus control (mock). The means ± SEM are shown from at least five independent experiments. (**C**) Primary MDMs from three donors overexpressing either CD169 or ACE2, or control were infected with S-pseudotyped lentivirus (20 ng p24^Gag^) for 3 days, followed by analysis of luciferase activity in whole cell lysates. (**D**) Untransduced primary MDMs (representative of 3 donors) were pre-treated with anti-CD169 mAb (20 μg/ml, 7D2), IgG1, or empty media for 30 min at 4°C prior to infection with S-pseudotyped lentivirus (20 ng p24^Gag^) for 3 days, followed by analysis of luciferase activity. RLUs from each donor in each group were normalized to no virus control (mock). The means ± SEM from at least 3 independent experiments are shown. *P*-values: paired t-test, two-tailed (**A**), one-way ANOVA followed by the Dunnett’s post-test (**B**) or Tukey’s post-test comparing to parental (THP1) cells (**C**) or each pre-treatment condition (D). ***: *p* < 0.001, ****: *p* < 0.0001, ns: not significant.

We next sought to determine the role of CD169 in mediating SARS-CoV-2 infection of macrophages. PMA-differentiated THP1 macrophages expressing wt or mutant CD169, ACE2, or CD169/ACE2 were infected with SARS-CoV-2 S-pseudotyped lentiviruses. Expression of wt CD169 on THP1/PMA macrophages enhanced S-pseudotyped lentiviral infection by ∼30-fold compared to parental THP1/PMA macrophages, similar to the levels of infection observed with ACE2^+^ THP1/PMA macrophages (**Fig. 2B**). In contrast expression of CD169/R116A did not enhance S-pseudotyped lentiviral infection of THP1/PMA cells, confirming that recognition of sialylated motifs on SARS-CoV-2 S is essential for CD169-mediated infection of macrophages (**Fig. 2B**). Co-expression of wt CD169 and ACE2 further enhanced S-pseudotyped lentiviral infection by greater than 11- fold and 3-fold when compared to cells expressing CD169 or ACE2, respectively (**Fig. 2B**), confirming that CD169 facilitates S-mediated entry into macrophages in the absence of ACE2 and enhances entry in the presence of ACE2. To confirm the role of CD169 in SARS-CoV-2 S-mediated infection in primary human MDMs, and given the relatively low and variable expression of endogenous CD169 in primary MDMs (**Fig. S3B**), we used lentiviral transduction to overexpress either wt CD169 or ACE2 (**Fig. S3C and D**). Following infection of transduced primary macrophages with S-pseudotyped lentivirus, we observed that CD169 or ACE2 overexpression in MDMs from multiple donors significantly enhanced SARS-CoV-2 S-pseudotyped lentivirus infection compared to control MDMs transduced with empty vector (**Fig. 2C**). Crucially, pre-treatment with anti-CD169 blocking mAb (7D2) prior to infection with S-pseudotyped lentivirus significantly attenuated infection of untransduced primary MDMs when compared to non-specific IgG1 pretreatment (**Fig. 2D**). These findings suggest that CD169 facilitates SARS-CoV-2 S-dependent fusion and entry into both THP1/PMA macrophages and primary MDMs in the absence of ACE2.

### SARS-CoV-2 establishes abortive infection in macrophages lacking ACE2

To investigate whether CD169 expression is sufficient to establish productive SARS-CoV- 2 infection and replication in ACE2-deficient macrophages, we infected THP1/PMA and primary MDMs overexpressing CD169, ACE2, or parental cells (lacking both CD169 and ACE2) with replication-competent SARS-CoV-2 (Washington isolate, NR-52281), as previously described (60). To evaluate productive infection, we examined the temporal expression of double-stranded RNA (dsRNA), a viral replication intermediate (61), as well as viral nucleocapsid (N) protein. SARS-CoV-2-infected THP1/PMA macrophages were fixed at various time points post infection and subjected to immunofluorescence analysis using antibodies against dsRNA and SARS-CoV-2 N. In contrast to parental THP1/PMA macrophages that showed background staining of dsRNA and no expression of SARS-CoV-2 N at any time point post infection (**Fig. 3A**), CD169-expressing THP1/PMA cells showed low levels of dsRNA production and small puncta staining of SARS-CoV-2 N that did not significantly increase over the course of infection (**Fig. 3B**). However, both ACE2^+^ (**Fig. 3C**) and CD169/ACE2 double-positive (**Fig. 3D**) THP1/PMA macrophages showed robust dsRNA and SARS-CoV-2 N production starting as early as 2-4 hours post infection (hpi), which significantly increased over the course of infection. There was a clear distinction in the spatial distribution of viral N protein in infected ACE2^+^ and CD169^+^/ACE2^+^ THP1/PMA macrophages over time, as noted by a transition from 6 hpi onward from small cytosolic N protein puncta to homogenous distribution throughout the cytosol (**Fig. 3C** and **D**). Furthermore, there was extensive co-localization between peri- nuclear dsRNA foci and N staining in ACE2^+^ and CD169^+^/ACE2^+^ THP1/PMA macrophages at 6 and 24 hpi. In accordance with our S-pseudotyped lentivirus infection data (**Fig. 2B**), co-expression of CD169 and ACE2 led to an increase in SARS-CoV-2 infection compared to ACE2-expressing cells, with higher levels and slightly earlier production of dsRNA and SARS-CoV-2 N (compare **Fig. 3C** and **D**). In contrast, N protein staining was not dispersed in the cytoplasm of CD169^+^ THP1/PMA macrophages and remained as small cytosolic puncta over the course of infection with minimal co- localization with dsRNA staining (**Fig. 3B**). These findings suggest that CD169-mediated SARS-CoV-2 entry in THP1/PMA macrophages results in initiation of viral transcription and moderate viral protein production.

**Figure 3.**
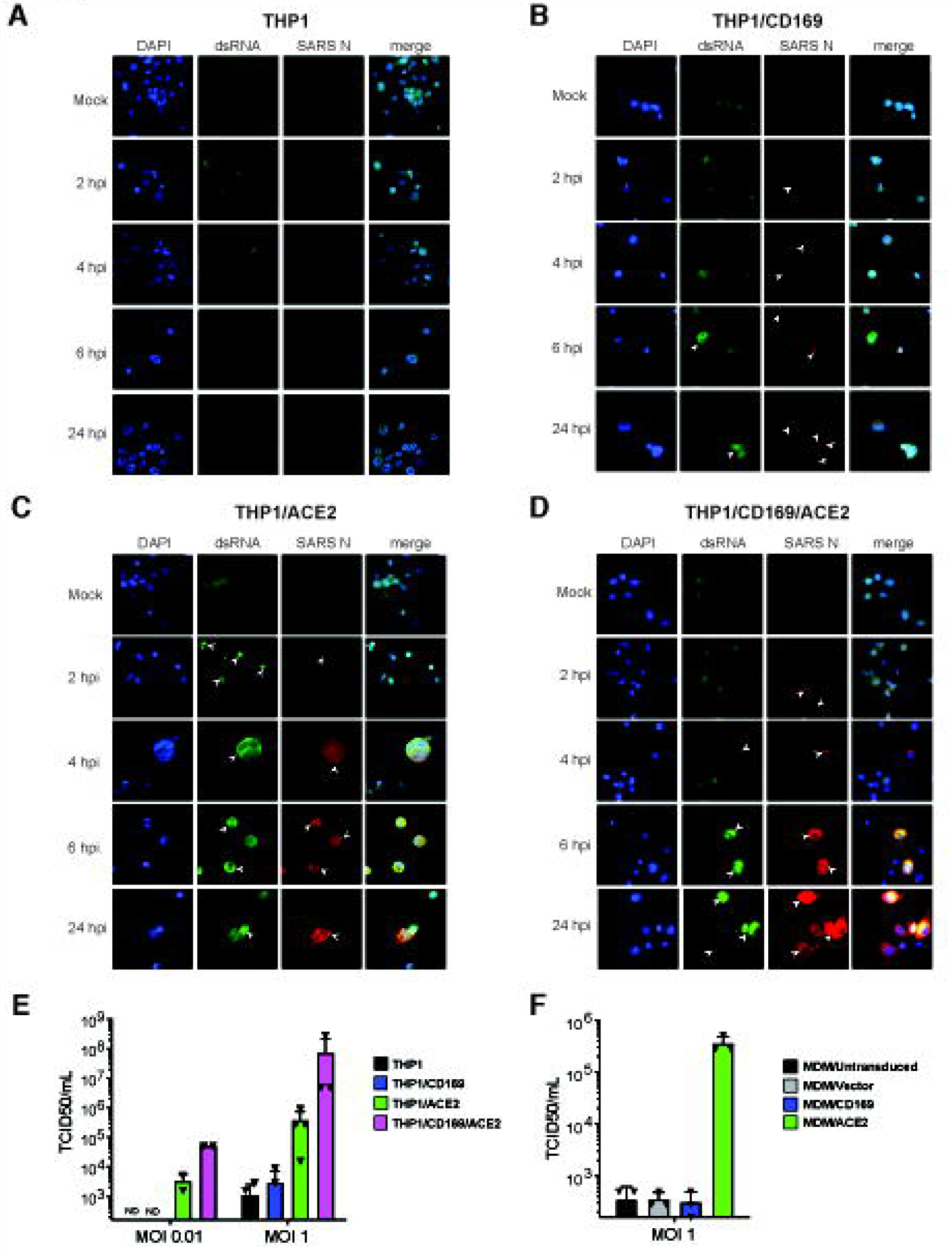
SARS-CoV-2 establishes restricted infection in CD169^+^ macrophages. (**A-D**) Representative immunofluorescence images (100x) of THP1/PMA macrophages infected with SARS-CoV-2 (MOI=1) and stained for nucleus (DAPI, blue), SARS-CoV-2 dsRNA (green), and SARS-CoV-2 nucleocapsid (red), at indicated timepoints post infection. Images shown for each THP1 cell line; untransduced control (parental, **A**), CD169-expressing (**B**), ACE2-expressing (**C**), and CD169/ACE2 double expressing (**D**). Bar=25 µm. (**E-F**) Culture supernatants from SARS-CoV-2 infected THP1/PMA cells (**E**) and primary MDMs (**F**) were harvested at 24 hpi and viral titers determined by TCID_50_ assay. The means ± SEM from at least 3 independent experiments are shown. LOD: limit of detection of assay.

Similar to our findings in THP1/PMA macrophages, we observed SARS-CoV-2 N expression in primary MDMs overexpressing ACE2, whereas we were not able to detect N protein in primary untransduced MDMs or those overexpressing CD169 at 24 hpi (**Fig. S4**). The lack of detection of either N (**Fig. S4**) or dsRNA (data not shown) in primary MDMs compared to the low level of N and dsRNA expression observed in THP1/CD169 macrophages (**Fig. 3B**) could be explained by the inefficient lentivector transduction efficiency and low expression of CD169 in primary MDMs (**Fig. S3B** and **C**) compared to the robust CD169 expression in retrovirally transduced THP1 cells (**Fig. S3A**). We further quantified infectious SARS-CoV-2 particle production from both THP1/PMA macrophages (**Fig. 3E**) and primary MDMs (**Fig. 3F**) by TCID50 assay and detected no infectious virions in culture supernatants from CD169-expressing cells. However, both THP1/PMA and primary MDMs expressing ACE2 showed robust production of infectious SARS-CoV-2 particles, with a marked increase in infectious virus production in THP1/PMA cells co-expressing CD169 and ACE2 compared to ACE2 alone (**Fig. 3E**). These results suggest an absence of post-entry restrictions to SARS-CoV-2 replication in macrophages, if virus entry is mediated by ACE2.

### CD169-mediated SARS-CoV-2 infection of macrophages results in *de novo* **expression of viral mRNAs**

To explore the spatial and temporal distribution of SARS-CoV-2 RNA at single cell level, single-molecule RNA FISH (smFISH) analysis (62) was performed in infected THP1/PMA macrophages. Individual fluorescent spots corresponding to viral gRNAs were detected in CD169^+^, ACE2^+^ and CD169^+^/ACE2^+^ THP1/PMA macrophages, as early as 1 hpi (**Fig. S5**). By 6 hpi, we observed diffuse cytosolic staining and perinuclearly localized bright gRNA foci (**Fig. S5**), which further increased in in ACE2^+^ and CD169^+^/ACE2^+^ but not CD169^+^ THP1/PMA macrophages (**Fig. S5**). In order to distinguish between gRNAs and N gene transcripts, we probed cells at 24 hpi with two probe sets labeled with distinguishable dyes one against SARS-CoV-2 ORF1a and the other against N gene. The former detects only gRNA and the later reports both gRNA and N sub-genomic transcripts **(Fig. 4A)**. The staining, particularly that of N gene, showed diffused cytoplasmic distribution, only in ACE2^+^ and CD169^+^/ACE2^+^ THP1/PMA macrophages, especially in the advanced stages of infection.

**Figure 4.**
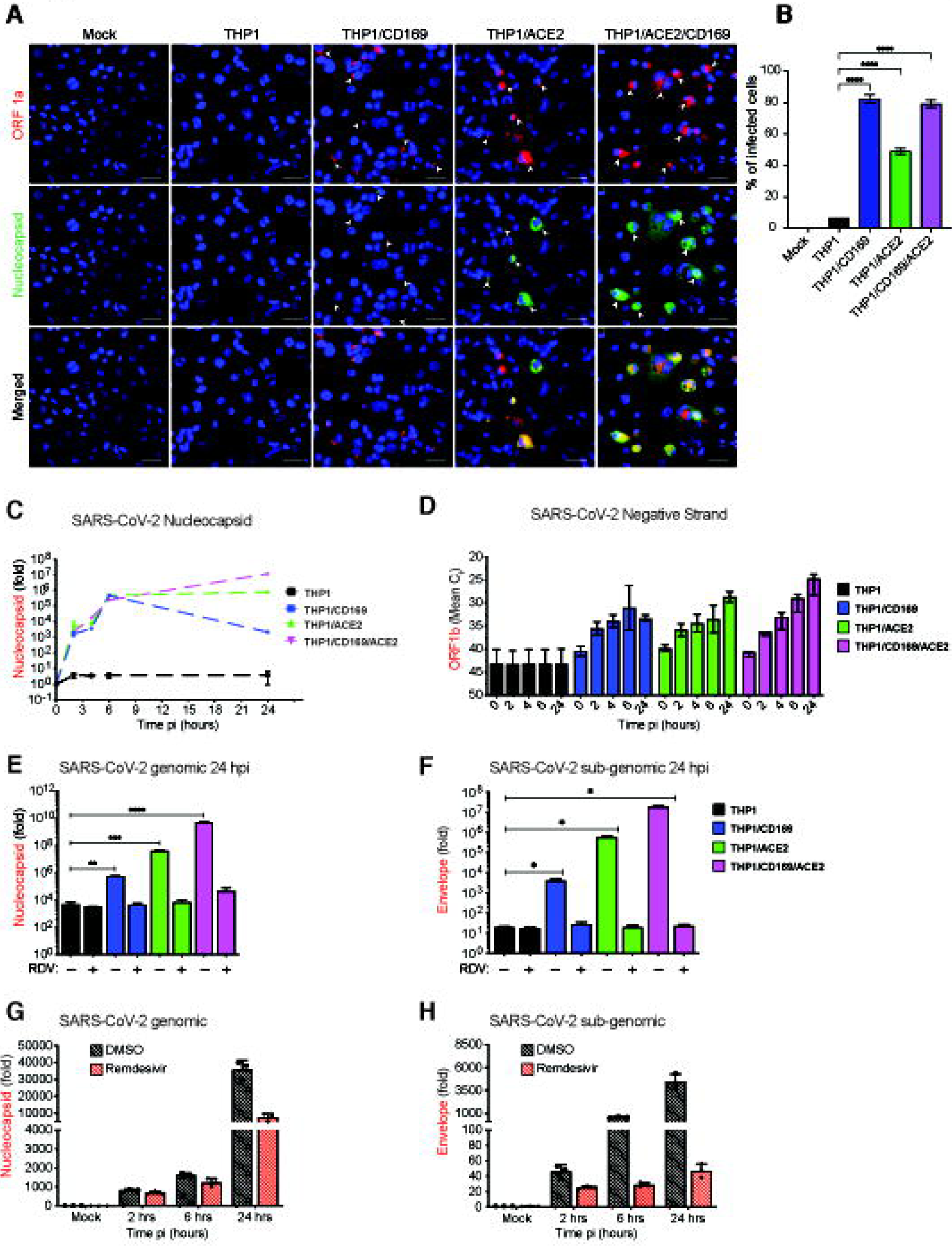
CD169-mediated SARS-CoV-2 infection of macrophages results in restricted viral RNA expression. (**A**) Single molecule RNA FISH analysis for viral RNAs using ORF1a specific probe set (labeled with Texas Red) and N specific probe set (labelled with TMR) in SARS-CoV-2 infected THP1/PMA macrophages (MOI=10, 24 hpi) (**B**) Percentage of infected cells based on the presence of SARS-CoV-2 RNAs determined from 10-20 independent fields, (representative field shown in **A**). Data are representative of 2 independent experiments. Bar=25 µm. (**C-F**) THP1/PMA macrophages infected with SARS-CoV-2 (MOI=1) in the absence (**C-D**) or presence (**E-F**) of remdesivir (RDV, 1 μM). Total RNA was harvested at indicated times post-infection and expression of viral transcripts was quantified by RT- qPCR. Replication kinetics of SARS-CoV-2 was quantified by (**C**) total N amplification (values from each group normalized to mock (uninfected) or (**D**) negative strand genome from strand-specific reverse-transcription and ORF1b amplification (mean Ct values for each condition). Expression of total N mRNA (**E**) and Envelope sgRNA (**F**) transcripts in THP1/PMA cells pretreated with RDV was analyzed at 24 hpi. Values from each group normalized to mock (uninfected). (**G-H**) MDMs from multiple donors were infected with SARS-CoV-2 (MOI=1), in the absence or presence of RDV (1 μM), and viral replication kinetics analyzed by RT-qPCR at indicated timepoints for total N mRNA (**G**), Envelope sgRNA (**H**) transcripts. The means ± SEM are shown from at least 3 independent experiments, each symbol represents a different donor. Significant differences between groups were determined by one-way ANOVA followed by the Dunnett’s post-test (**E-F**), comparing to control parental THP1 cells. *P*-values: *<0.1; **<0.01; ***<0.001; ****<0.0001.

Dispersion of N transcripts (gRNA and sgRNA) in the cytosol of ACE2^+^ and CD169^+^/ACE2^+^ THP1/PMA macrophages mirrors the temporal localization phenotype of viral N protein (**Fig. 3C** and **D**), and might be indicative of formation of N positive viral replication compartments (63). While a majority of virus-exposed CD169^+^ THP1/MA macrophages expressed fluorescent puncta (gRNA) at 24 hpi (indicated by the white arrowheads showing both ORF1a and N expression, **Fig. 4A and B**), transition to bright gRNA foci or cytosolic expansion was not observed, suggesting that CD169 expression in THP1/PMA cells increased uptake of virus, but without ACE2, failed to establish viral replication foci. It should be noted that the formation of distinct fluorescent puncta in CD169^+^ THP1/PMA macrophages at 24 hpi is suggestive of *de novo* RNA synthesis and not virus inoculum, since these puncta were not detected at 1 hpi (**Fig. S5**). While viral RNAs in the CD169-expressing cells are localized in few distinct granulated puncta, ACE2-expressing cells exhibited inclusion-like structures, suggesting that differential engagement of viral entry receptors (such as CD169 and ACE2) leads to altered fate of the incoming viral genome and subsequent steps in the viral replication cycle.

To further investigate the step at which SARS-CoV-2 replication is restricted in macrophages, THP1/PMA macrophages or those expressing either CD169, ACE2, or both CD169 and ACE2 were infected with SARS-CoV-2 and cells lysed and harvested at 2, 4, 6, and 24 hpi to quantify the level of total SARS-CoV-2 N transcripts, +gRNA and sgRNA, as well as negative sense antigenomic RNA, by RT-qPCR. We detected increasing levels of total SARS-CoV-2 N transcripts at early time points (2-6 hpi) in THP1/PMA macrophages expressing CD169, ACE2, or both CD169 and ACE2, whereas parental THP1/PMA macrophages showed no significant increase in viral N RNA levels compared to mock infection controls (**Fig. 4C**). Furthermore, there were no significant differences in gRNA levels at early times post infection (up to 6 hpi) between CD169^+^, ACE2^+^ or CD169^+^/ACE2^+^ THP1/PMA macrophages suggesting an absence of cell-intrinsic restrictions to early steps of SARS-CoV-2 replication, such as attachment and fusion, in CD169+ THP1/PMA macrophages. Interestingly, SARS-CoV-2 N transcripts peaked at 6 hpi in CD169+ THP1/PMA cells, followed by a progressive decline at 24 hpi (**Fig. 4C**). In contrast, there was a temporal increase in SARS-CoV-2 N transcripts in cells expressing ACE2, with markedly higher expression at 24 hpi compared to CD169- expressing cells. In concordance with SARS-CoV-2 N protein expression (**Fig. 3D**),

THP1/PMA cells expressing both CD169 and ACE2 expressed the highest levels of SARS-CoV-2 N transcripts at 24 hpi compared to those expressing ACE2 or CD169 alone. After virus entry and capsid uncoating, the next steps in the SARS-CoV-2 replication cycle are translation of the viral gRNA followed by the formation of viral replication-transcription complexes, which enables synthesis of the negative sense antigenomic RNA (64). Negative sense antigenomic RNAs are present at significantly lower levels than +gRNA and sgRNA, and are templates for synthesis of additional positive sense gRNA and sgRNAs (64). We employed strand-specific RT-qPCR analysis to detect negative sense viral RNA (ORF1b) and confirmed viral replication (at 2 hpi) in CD169^+^ THP1/PMA cells, compared to complete absence of negative sense viral RNAs in parental THP1/PMA macrophages (**Fig. 4D**). Interestingly, negative sense RNA expression in CD169^+^ THP1/PMA macrophages plateaued at 6 hpi, suggesting that CD169-mediated SARS-CoV-2 entry promotes initial steps of viral replication. In contrast, there was a progressive increase in negative sense RNA levels in ACE2^+^ THP1/PMA macrophages particularly at 24 hpi, and a further enhancement was observed upon co-expression of CD169 and ACE2 in THP1/PMA macrophages at later times post infection (**Fig. 4D**).

To confirm that temporal increases in viral transcripts in CD169^+^ or ACE2^+^ THP1/PMA macrophages were generated from ongoing virus transcription, THP1/PMA cells (±CD169 ±ACE2) were pre-treated with remdesivir (RDV), a known inhibitor of SARS-CoV-2 RNA synthesis, prior to infection with SARS-CoV-2 (24, 63). RT-qPCR analysis of both gRNA (N transcripts) (**Fig. 4E**) and sgRNA (E transcripts) (**Fig. 4F**) transcripts harvested at 24 hpi revealed that RDV pre-treatment completely blocked the increase in viral RNA expression in THP1/PMA macrophages expressing CD169, ACE2, or both CD169 and ACE2, to levels observed in parental THP1/PMA macrophages. Furthermore, levels of viral gRNA (**Fig. 4G**) and sgRNA (**Fig. 4H**) in SARS-CoV-2 infected untransduced MDMs progressively increased over time, and these increases in viral gRNA and sgRNAs were also suppressed upon RDV pre-treatment. Notably, primary MDMs harbored higher levels of viral transcripts compared to parental THP1/PMA macrophages at all times post infection (**Fig. 4E-H**), suggesting that endogenous CD169 expression was sufficient to facilitate entry and initiation of virus replication in primary MDMs. The marked lower levels of viral transcripts in primary MDMs (**Fig. 4G** and **H**) compared to CD169^+^ THP1/PMA macrophages (**Fig. 4E** and **F**) might be correlated to the lower CD169 expression in primary MDMs compared to CD169^+^ THP1/PMA macrophages cells which were selected for high CD169 expression (**Fig. S3A**). Taken together, these results suggest that CD169-mediated viral entry enables initiation of viral RNA replication and transcription in ACE2-deficient CD169^+^ macrophages even in the absence of establishment of robust viral RNA replication foci in macrophages, and that SARS-CoV-2 infection of macrophages is restricted at a post-entry step of virus replication.

### Low level expression of SARS-CoV-2 genomic and sub-genomic RNAs is sufficient to induce pro-inflammatory cytokines in non-productively infected macrophages

Since CD169 expression in macrophages was sufficient to permit SARS-CoV-2 entry and initiate restricted viral RNA expression, we investigated whether *de novo* viral RNA synthesis in the absence of productive virus replication can trigger innate immune responses. SARS-CoV-2 infected THP1/PMA cells (±CD169 ±ACE2) were lysed at 24 hpi, and total RNA was analyzed by RT-qPCR for inflammatory cytokine expression (**Fig. 5**). We observed significant induction of expression of IL-6, TNFα, IL-1β, and IL-18 mRNA in non-productively infected CD169^+^ THP1/PMA macrophages compared to the parental THP1/PMA cells (**Fig. 5A**). IL-6 and TNFα mRNA expression in SARS-CoV-2 infected THP1/CD169 cells was similar to the levels observed in productively infected THP1/ACE2, but lower than that observed in THP1/CD169/ACE2 cells. Intriguingly, induction of pro-inflammatory cytokines, IL-1β and IL-18, was only consistently observed in non-productively infected THP1/CD169 macrophages but not in the productively infected THP1/ACE2 or THP1/CD169/ACE2 macrophages (**Fig. 5A**). A delayed or impaired type I IFN response is thought to be a critical mechanism for COVID-19 pathogenesis, though impaired induction of type I IFN response has mostly been reported in infected lung epithelial cells (65). Interestingly, we observed muted induction of IFNβ, IP-10 and Viperin in SARS-CoV-2 infected CD169^+^ THP1/PMA macrophages. In contrast, IFNβ, IFNλ1, IP- 10 and Viperin mRNA expression was dramatically upregulated upon establishment of productive SARS-CoV-2 infection in THP1/ACE2 and THP1/CD169/ACE2 macrophages (**Fig. 5A**) and correlated with the extent of viral RNA expression and virus replication in these cells (**Fig. 3**).

**Figure 5.**
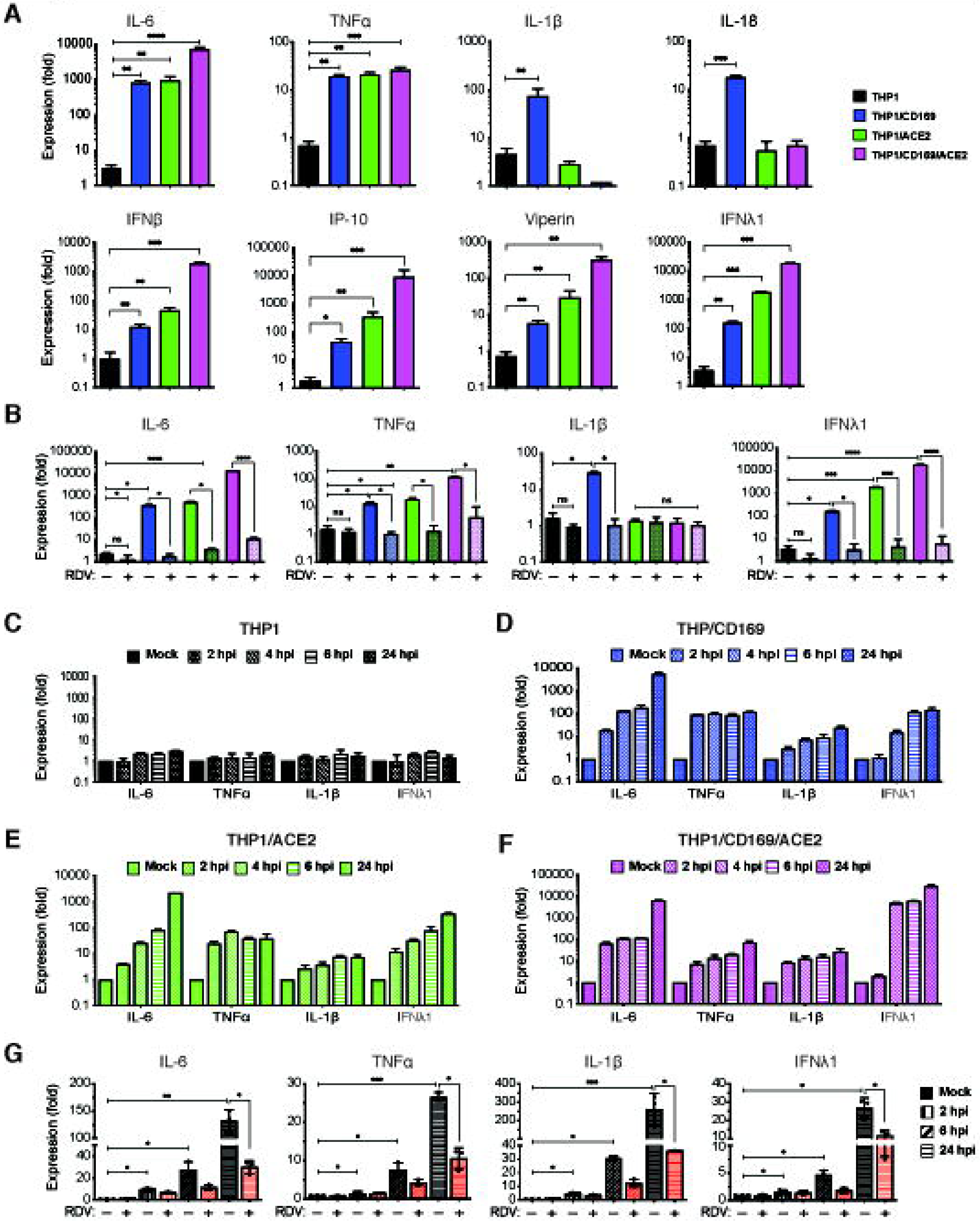
Restricted expression of SARS-CoV-2 gRNA and sgRNA induces pro-inflammatory responses in non-productively infected macrophages. (**A-B**) PMA-differentiated THP1 cells from parental (THP1) or those expressing CD169, ACE2, or CD169/ACE2 were infected with SARS-CoV-2 (MOI=1, 24 hpi) in the absence or presence of RDV (1 μM, 30 mins), and total RNA was quantified by RT-qPCR for pro-inflammatory cytokines and ISGs. Values were normalized to mock-infected control for each group. Fold-induction for indicated pro-inflammatory cytokines (**A**, top panel) and ISGs (**A**, bottom panel) in the absence of RDV, or in the presence of RDV (**B**). (**C-F**) Kinetics of pro-inflammatory cytokine/ISG mRNA expression in the absence of RDV in parental (**C**), CD169-expressing (**D**), ACE2-expressing (**E**), and CD169/ACE2-expressing (**F**) PMA-differentiated THP1 cells. (**G**) MDMs from multiple donors were infected with SARS-CoV-2 (MOI=1, 24 hpi) and total RNA analyzed by RT-qPCR. MDMs were pre-treated with DMSO (control) or RDV (1 μM, 30 mins) prior to infection, and fold-induction of indicated cytokine mRNAs normalized to mock-infected controls from each donor. RDV: Remdesivir, ISGs: interferon-stimulated genes. Minus sign represents DMSO, and plus sign represents RDV pre-treatment. The means ± SEM from at least 3 independent experiments are shown. Significant differences between groups were determined by one-way ANOVA followed by the Dunnett’s post-test between groups (**A**) or Tukey’s multiple comparisons test within groups (**B** and **G**), comparing to parental THP1 (**A**) or DMSO-treated (**B**, **G**) in each group. *P*-values: *<0.1; **<0.01; ***<0.001; ****<0.0001, ns: not significant.

Since previous reports have suggested that macrophage exposure to recombinant S protein may trigger inflammatory cytokine production (17, 66), we infected parental, CD169^+^, ACE2^+^, or CD169^+^/ACE2^+^ THP1/PMA cells with S-pseudotyped lentivirus but observed no induction of pro-inflammatory cytokines (IL-6, TNFα, IL-1β, and IFNλ1) (**Fig. S6**). To confirm that the induction of pro-inflammatory cytokine and ISG expression was due to viral replication and sensing of viral RNA, we infected THP1/PMA macrophages with SARS-CoV-2 in the presence of RDV. Pre-treatment with RDV which abolished virus replication (**Fig. 4E** and **F**) completely abrogated IL-6, TNFα, IL-1β and IFNλ1 induction in CD169^+^, ACE2^+^, and CD169^+^/ACE2^+^ THP1/PMA macrophages (**Fig. 5B**), suggesting that *de novo* viral RNA transcription and sensing of virus replication intermediates are required for the induction of pro-inflammatory cytokines and ISGs in macrophages. Temporal analysis of inflammatory cytokine induction revealed that while SARS-CoV-2 infection of parental THP1/PMA cells did not result in induction of pro-inflammatory responses (**Fig. 5C**), infection of CD169^+^ THP1/PMA cells led to rapid induction of IL-6, TNFα, IL-1β, and IFNλ1 mRNA expression (**Fig. 5D**). In fact, fold-induction in levels of pro-inflammatory cytokines at early times post infection (4-6 hpi) was similar between CD169^+^ and ACE2^+^ macrophages (**Fig. 5D** and **E**), indicating that early viral RNA production is the key trigger of innate immune activation. Accelerated kinetics and highest magnitude of induced IFNλ1 expression was observed in SARS-CoV-2 infected CD169^+^/ACE2^+^ THP1/PMA macrophages (**Fig. 5F,** note difference in scale on y axis), which correlated with greater level of negative sense antigenomic RNA (**Fig. 4D**) and highest levels of virus replication in CD169^+^/ACE2^+^ THP1/PMA macrophages (**Fig. 3E**).

To further confirm the findings obtained from CD169^+^ THP1/PMA macrophages, we pre-treated primary MDMs from multiple donors with RDV for 30 minutes prior to infection with SARS-CoV-2. Total RNA was harvested and subsequently analyzed by RT-qPCR to determine fold-induction of IL-6, TNFα, IL-1β, and IFNλ1 mRNA expression at indicated timepoints (**Fig. 5G**). Similar to the observations in CD169^+^ THP1/PMA macrophages (**Fig. 5B**), SARS-CoV-2 infection induced IL-6, TNFα, IL-1β, and IFNλ1 mRNA in primary MDMs (**Fig. 5G**), with proinflammatory cytokine induction observed as early as 2 hpi. Furthermore, induction of pro-inflammatory responses in primary MDMs was significantly suppressed by RDV pre-treatment (**Fig. 5G**), confirming that establishment of infection and initiation of viral transcription is required for macrophage-intrinsic innate immune activation. Collectively, these findings suggest that CD169-mediated restricted SARS-CoV-2 infection of macrophages induces robust inflammatory responses.

### Cytosolic RNA sensing by RIG-I and MDA-5 is required for SARS-CoV-2 induced inflammation in macrophages

Since innate immune activation in THP1/CD169 cells and primary MDMs requires the expression of *de novo* viral RNAs (**Fig. 5B** and **G**), we next sought to delineate the nucleic acid sensing mechanism required for the detection of SARS-CoV-2 gRNA and sgRNAs in CD169^+^ THP1/PMA macrophages. Depending on the specific pathogen-derived cytosolic nucleic acids, numerous host sensors can detect and trigger innate immune activation via the MAVS and/or STING pathways (67). Viral RNAs can be sensed by RIG-I-like receptors (RLRs) or endosomal toll-like receptors (TLRs) to activate MAVS or TRIF, respectively, leading to induction of pro-inflammatory cytokines (68). Innate immune sensing of SARS-CoV-2 RNAs by RLRs such as RIG-I and MDA-5, or endosomal TLRs (TLR7/8), has been previously proposed (69, 70). It has also been hypothesized that SARS-CoV-2-induced mitochondrial damage and release of mitochondrial DNA into the cytosol, a cellular stress response, could trigger cGAS/STING sensing pathway in infected cells (71, 72). To investigate which of the nucleic acid sensing pathways are involved in sensing abortive SARS-CoV-2 infection in macrophages, we stably knocked-down expression of either RIG-I, MDA-5, UNC93B1 (a protein required for TLR3/7/9 trafficking to endosomes, (73)), MAVS or STING in CD169^+^ THP1/PMA cells through shRNA-based lentiviral transduction. Upon successful selection of transduced cells targeted for knockdown of individual host proteins, we observed robust decrease in both mRNA (**Fig. 6A**) and protein (**Fig. 6B**) expression. Knock-down of RIG-I, MDA-5, UNC93B1, MAVS or STING in THP1/CD169 macrophages did not impact infection efficiency of SARS-CoV-2, as shown by RT-qPCR analysis of viral gRNA and sgRNAs in each cell line (**Fig. 6C** and **D**). While knockdown of UNC93B1 in virus-infected THP1/PMA CD169^+^ cells had negligible impact on induction of pro-inflammatory cytokines (IL-6, TNFα, IL-1β and IFNλ1) compared to scramble control cells, depletion of either RIG-I or MDA-5 led to dramatic reduction in pro-inflammatory cytokine expression to near background levels, suggesting both cytosolic viral RNA sensors, but not endosomal TLRs, are required for innate immune sensing of SARS-CoV- 2 transcripts (**Fig. 6E-H**). Furthermore, knock-down of MAVS but not STING, completely abrogated SARS-CoV-2- induced innate immune activation in THP1/PMA macrophages, further confirming the requirement of cytosolic viral RNA sensing for induction of pro-inflammatory cytokines.

**Figure 6.**
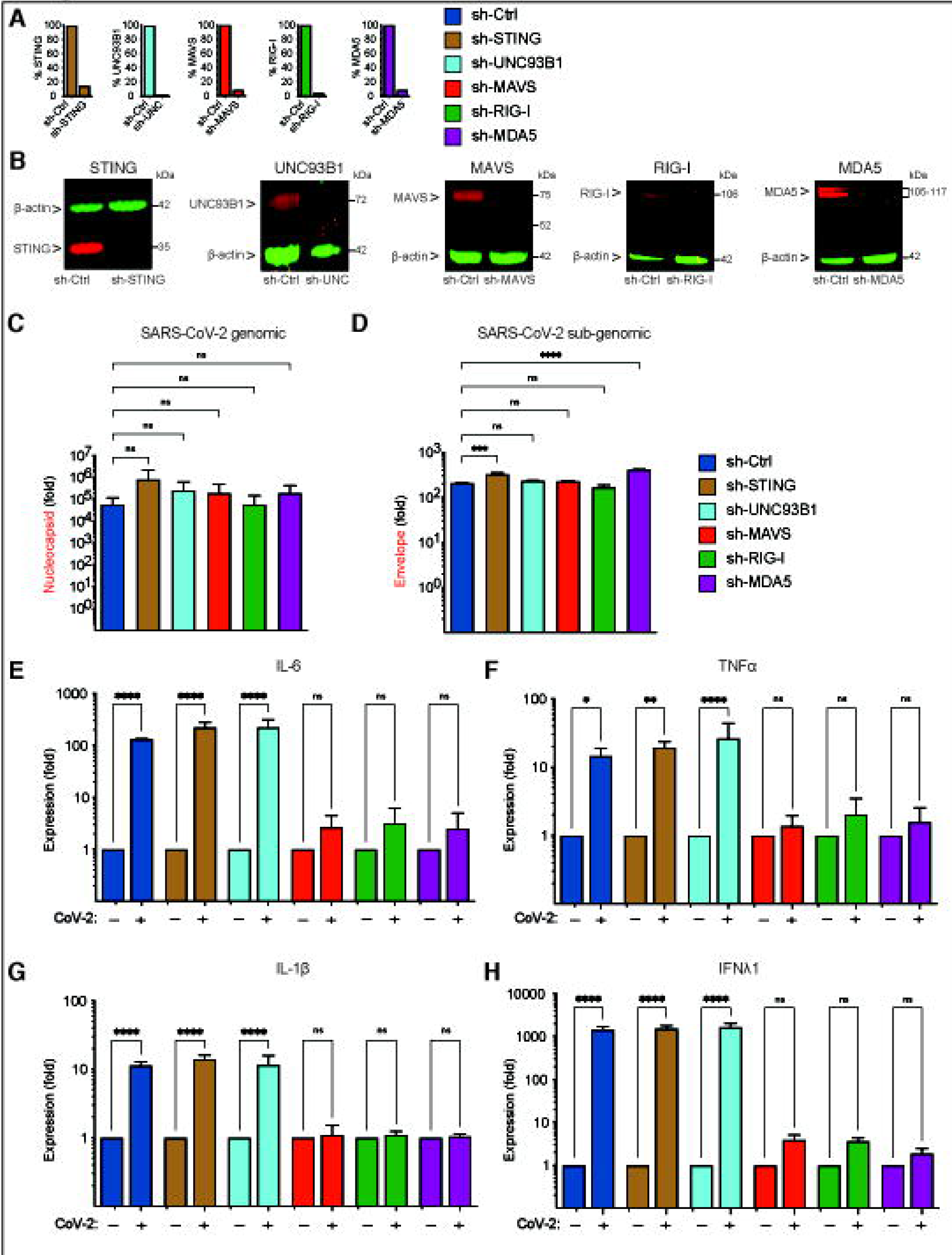
Cytosolic RNA sensing by RIG-I and MDA-5 is required for SARS-CoV-2 induced innate immune responses in macrophages. (**A-B**) CD169-expressing THP1 monocytes were transduced with lentivectors expressing shRNA against non-specific control (scramble), or specific sequences against STING, UNC93B1, MAVS, RIG-I, and MDA-5. Knockdown of host proteins targeted by shRNAs compared to scramble analyzed by RT-qPCR (**A**), and immune blotting (**B**). (**C-H**) CD169^+^ THP1 cells with stable knockdown of target genes were PMA-differentiated and infected with SARS-CoV-2 (MOI=1, 24 hpi), and total RNA was analyzed by RT-qPCR for total nucleocapsid (gRNA) (**C**), sgRNA (Envelope) (**D**), and pro-inflammatory cytokines/ISGs, IL6, TNFα, IL-1β and IFNλ1 (**E-F**), normalized to mock-infected controls for each group. Each knockdown is represented by different colors as in **A**. The means ± SEM from 3 independent experiments are shown, and significant differences were determined by one-way ANOVA followed by the Dunnett’s post-test comparing to scramble THP1 (**C**-**D**), or by two-way ANOVA followed by Bonferroni post-test comparing Mock to CoV-2^+^ in each group (**E**-**H**). *P*-values: *<0.1; **<0.01; ***<0.001; ****<0.0001), ns: not significant.

### MAVS is essential for SARS-CoV-2 RNA-induced innate immune activation in both non-productively and productively infected THP1/PMA macrophages

We next sought to determine if MAVS was required for induction for pro-inflammatory responses not only in CD169-mediated abortive infection but also in ACE2-mediated productive infection of macrophages with SARS-CoV-2. Parental THP1 cells or those expressing CD169, ACE2, or both CD169 and ACE2 were transduced with lentivectors expressing scramble shRNAs (control) or MAVS-specific shRNAs, which led to robust knock-down of MAVS protein expression in all cell lines (**Fig. 7A**). While MAVS depletion did not affect subsequent SARS-CoV-2 infection of CD169^+^, ACE2^+^ or CD169^+^/ACE2^+^ THP1/PMA cells, as quantified by total SARS-CoV-2 N gRNA transcripts (**Fig. 7B**), induction of pro-inflammatory cytokines (IL-6, TNFα, IL-1β and IFNλ1) was significantly attenuated in both productively (ACE2^+^, CD169^+^/ACE2^+^) and abortively (CD169^+^) infected THP1/PMA macrophages upon MAVS knockdown (**Fig. 7C**). These results suggest that MAVS plays a pivotal role in pro-inflammatory cytokine induction in both ACE2-dependent and independent (CD169-mediated) SARS-CoV-2 infection of THP1/PMA macrophages. Taken together, these findings suggest that CD169-dependent establishment of abortive SARS-CoV-2 infection in macrophages triggers RIG-I/MDA-5 mediated sensing of viral gRNA and sgRNAs and MAVS-dependent inflammatory cytokines induction, which might contribute to the dysregulated hyper-immune phenotype of inflammatory macrophages and severity of COVID-19 disease.

**Figure 7.**
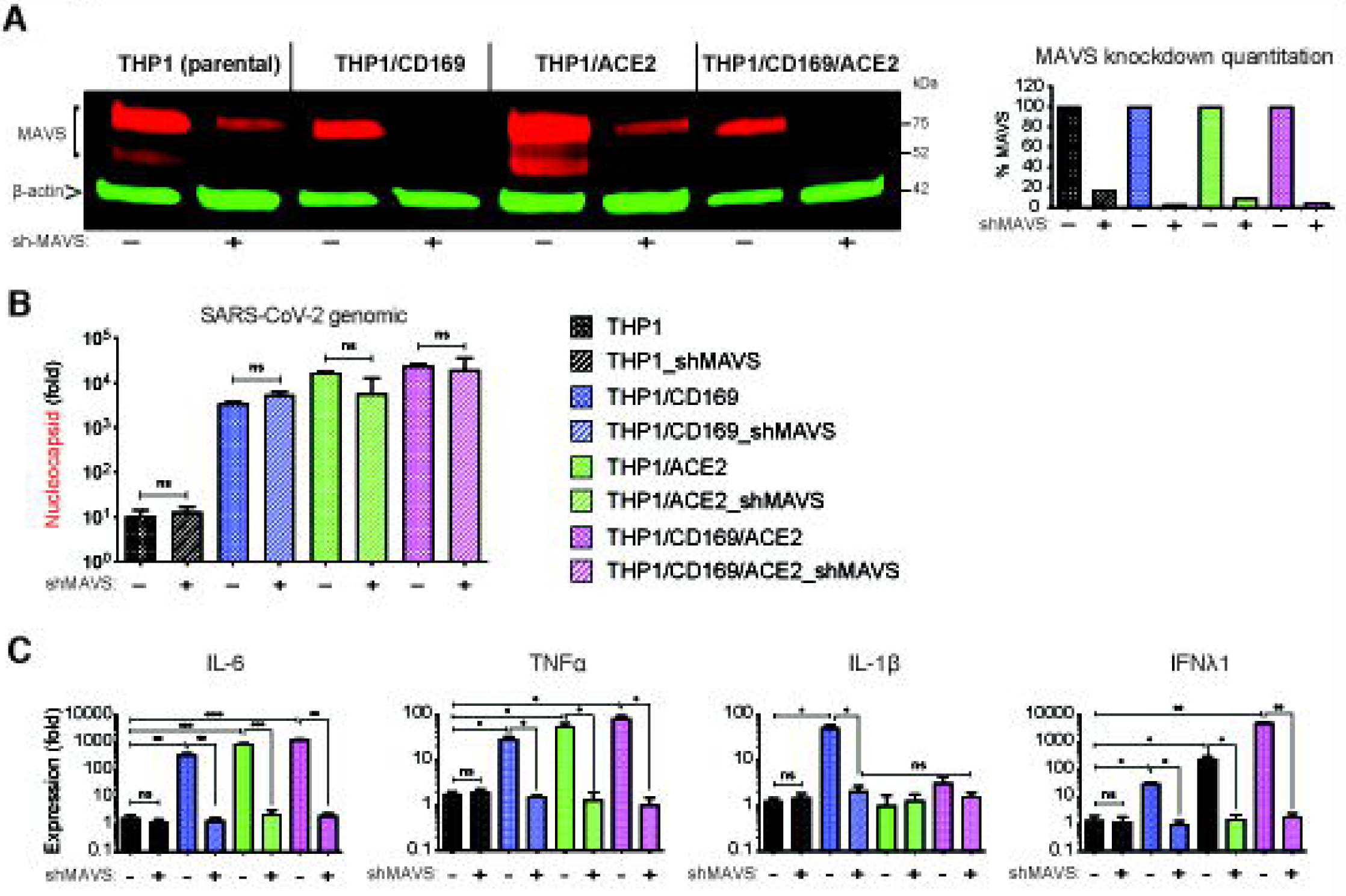
MAVS is essential for SARS-CoV-2 RNA-induced inflammatory responses in macrophages. (**A**) Parental (THP1) monocytes and those expressing CD169, ACE2, or CD169/ACE2 were transduced with lentivectors expressing shRNA against scrambled sequence (-), or MAVS sequence (+), and knockdown of MAVS in each cell line analyzed by immune blotting (**A**, left panel), and quantified (**A**, right panel). (**B-C**) THP1 monocytes with stable MAVS knockdown were PMA-differentiated and infected with SARS-CoV-2 (MOI=1, 24 hpi), and total RNA analyzed by RT-qPCR for (**B**) total viral transcripts (nucleocapsid), and (**C**) IL6, TNFα, IL-1β and IFNλ1 mRNA expression, normalized to mock-infected controls for each group. The means ± SEM from 3 independent experiments are shown, and significant differences were determined by one-way ANOVA followed by the Tukey-Kramer post-test within groups (**B**, and **C**), or Dunnett’s post-test between groups (**C**). *P*- values: *<0.05; **<0.01; ***<0.001; ****<0.0001, ns: not significant.

## Discussion

Though monocytes and macrophages have not been directly implicated in productive SARS-CoV-2 infection, several reports have provided evidence of SARS-CoV-2 RNA and antigen in circulating monocytes, macrophages, and tissue-resident alveolar macrophages, although these cells are not known to express ACE2 (**Fig. 1** and (30). Importantly, macrophage-intrinsic inflammatory phenotype in lung BALF samples has been associated with COVID-19 disease severity (7). In this study, we examined the mechanisms by which macrophages potentially contribute to inflammation during SARS-CoV-2 infection, using THP1/PMA and primary human macrophages expressing a myeloid cell-specific receptor, CD169, in the presence or absence of ACE2. We showed that CD169 expression mediated entry and fusion of SARS-CoV-2 S-pseudotyped lentivirus in both CD169^+^ THP1/PMA and primary macrophages. It was recently reported that CD169 expressed on dendritic cells captures SARS-CoV-2 particles via binding to gangliosides exposed on the viral membrane, such as GM1, and mediates trans infection of bystander cells (19). Our results, however, suggest that CD169 can also bind the sialylated SARS-CoV-2 S protein. Other studies, including ours, have also described glycan-dependent interactions of SARS-CoV-2 S protein with C-type lectin receptors (DC-SIGN, LSIGN) and Tweety family member 2 (TTYH2) (17, 19, 74). While ACE2- independent mechanisms of SARS-CoV-2 interactions with myeloid cells can be mediated by diverse virus particle-associated antigens (gangliosides, mannosylated S and sialylated S), our results suggest that CD169-S interaction is sufficient to promote virus entry and fusion in macrophages in an ACE2-independent manner.

Pre-treatment with antibodies targeting SARS-CoV-2 S NTD and inhibition of endosomal cathepsins markedly attenuated S-pseudotyped lentiviral infection in CD169^+^ macrophages, suggesting that S interaction with CD169 and virus entry mechanisms are distinct to those targeted by ACE2. This is in agreement with recent data suggesting that binding epitopes for myeloid cell-specific receptors are found outside the ACE2 binding domain in the spike receptor binding domain (RBD) (17). Consequently, the results presented here have implications for antibody treatment in COVID-19 patients, as current therapies are primarily focused on RBD-binding antibodies (57, 75–78). Targeting non-RBD epitopes to disrupt ACE2-independent entry in CD169^+^ myeloid cells could serve as a potential mechanism for preventing myeloid cell-intrinsic immune activation. Indeed, broadly neutralizing nanobodies and potent NTD-targeting neutralizing antibodies from COVID-19 patients that block both ACE2-dependent and ACE2-independent entry were recently described (57, 75–78) and might provide potential benefit in also suppressing macrophage-intrinsic inflammatory signature.

Efficiency of CD169-mediated S-pseudotyped infection in macrophages was similar to that mediated by ACE2 (**Fig. 2**). Importantly, CD169-mediated SARS-CoV-2 entry was also similarly efficacious to that mediated by ACE2 in THP1/PMA macrophages, as evident by the similar number of viral gRNA copies at 6 hpi (**Fig. 4C**), though viral gRNA and sgRNA expression in THP1/CD169+ macrophages was attenuated at later times post infection (**Fig. 4E** and **F**). SARS-CoV-2 RNA synthesis occurs within ER-derived double-membrane vesicles (DMVs). The establishment of viral replication factories within DMVs in the cytoplasm of infected cells is induced by viral proteins, in concert with cellular factors (64). Previous reports have utilized dsRNA staining to visualize these viral replication organelles (79). While viral dsRNA+ cells were observed in a minor percentage of THP1/CD169^+^ macrophages at 6 hpi (**Fig. 3B**), no further increase was observed. In contrast, majority of the virus-exposed THP1/ACE2 and THP1/CD169/ACE2 macrophages were dsRNA+ by 24 hpi (**Fig. 3C** and **D**). Since this staining strategy requires expression of high levels of viral dsRNA, the paucity of dsRNA positivity in SARS-CoV-2 infected CD169^+^ macrophages might reflect selective impairment of formation of DMVs. However, expression of ACE2 in THP-1/PMA and primary MDMs restored infectious virus particle production, suggesting that macrophages are permissive to SARS-CoV-2 replication when entry is facilitated by ACE2. Considering that both CD169 and ACE2 mediated virus entry resulted in similar levels of SARS-CoV- 2 gRNA at 6 h pi but only ACE2-mediated virus entry resulted in productive virus infection, these results implicate a hitherto unappreciated post-entry role for ACE2 in virus life cycle in macrophages. Interestingly, expression of both CD169 and ACE2 in macrophages led to enhanced kinetics and magnitude of infection, reflecting an entry-enhancing effect of CD169 even in the context of ACE2-mediated infection, though the mechanism of enhanced kinetics of SARS-CoV-2 replication in CD169^+^/ACE2^+^ macrophages remains unclear.

Despite lack of productive infection, cytoplasmic viral RNA expression in CD169^+^ macrophages potently induced expression of pro-inflammatory cytokines and chemokines. Thus, CD169-mediated viral entry does not simply enable viral uptake by macrophages but also initiates SARS-CoV-2 replication and triggers inflammatory cytokine expression. Critically, pre-treatment with RDV not only blocked *de novo* viral RNA expression but also significantly reduced pro-inflammatory cytokine expression in non-productively infected CD169^+^ macrophages (**Fig. 5**), suggesting that neither S protein interaction with cell surface receptors nor TLR-mediated sensing of incoming viral genome in the endosomal lumen is sufficient to trigger robust innate immune activation. Rather newly synthesized viral RNAs (negative sense gRNA, dsRNA, and/or sgRNAs) are the key drivers of innate immune activation in non-productively infected macrophages. While mRNA expression of type I and III IFNs peaked at 24 hpi and correlated with the extent of virus replication (marked by high levels of IFNλ1 mRNA expression at 24 hpi, **Fig. 5A**), pro-inflammatory cytokines, IL-6 and TNFα mRNAs were induced to comparable levels in both productively (ACE2^+^ and CD169^+^/ACE2^+^) and non-productively (CD169^+^) infected macrophages (**Fig. 5A**). Interestingly, significant induction of IL-1β and IL-18 was only observed in CD169+ THP1/PMA and primary macrophages (**Fig. 5D** and **G**), suggesting that CD169-mediated virus entry and infection establishment uncouples induction of inflammatory responses from robust viral replication. Since transcriptional priming of the inflammasome components, IL-1β and IL-18, was only observed in abortively infected CD169^+^ THP1/PMA macrophages (**Fig. 5A**), SARS-CoV-2 infected and primed macrophages might thus uniquely contribute to the inflammasome activation upon delivery of secondary activation signals and perpetuate the hyper-inflammatory inflammatory phenotype.

Previous reports have implicated both RIG-I and MDA-5 in cytosolic sensing of SARS-CoV-2 RNAs (80, 81), although the primary RNA sensor might be cell-type dependent. For instance, recent studies have implicated MDA-5 as the primary viral RNA sensor in lung epithelial cells (82), while other studies have found both MDA-5 and RIG-I sense SARS-CoV-2 infection in Calu-3 cells (81, 83). Our results implicate both RIG-I and MDA-5 in SARS-CoV-2 RNA sensing in macrophages, as knockdown of either RIG-I or MDA-5 significantly attenuated inflammatory cytokine induction in CD169^+^ macrophages. Despite previous studies suggesting that induction of IL-6 and TNFα might be MAVS-independent (84), knock-down of MAVS abrogated viral RNA sensing and induction of NF-κB-dependent inflammatory cytokines (IL-6, TNFα, IL-1β, IL-18) in productively infected (ACE2^+^) and abortively infected (CD169^+^) macrophages. Thus, we propose a co-sensing requirement of both RIG-I and MDA-5 for detecting viral replication intermediates and a pivotal role of MAVS in signal transduction in non-productively infected CD169^+^ macrophages. Intriguingly, induction of RIG-I/MDA-5/MAVS-dependent IFNβ and ISG expression was muted compared to the robust upregulation of NF-kB-dependent pro-inflammatory cytokines in SARS-CoV-2 infected CD169^+^ macrophages. Since RLR relocalization to the mitochondrial, peroxisomal or ER membranes is thought to initiate robust MAVS-dependent IRF3 activation (85–87), it is tempting to speculate that during restricted SARS-CoV-2 infection and diminished expression of viral RNAs, cytosolic retention or altered intracellular localization of RLRs might control the strength and specificity of downstream responses and favor NF-kB-dependent proinflammatory cytokine expression. Future studies will need to address spatiotemporal dynamics of RIG-I/MDA-5/MAVS interactions upon SARS-CoV-2 RNA sensing in macrophages.

SARS-CoV-2 RNA infection of lung epithelial cells can contribute to innate immune activation, inflammation, recruitment of inflammatory monocytes and macrophages to the alveolar space, and activation of tissue-resident alveolar macrophages (80, 81). Alveolar macrophages and airway epithelial cell-associated macrophages, which are uniquely positioned as gatekeepers to intercept invading pathogens from the bronchiolar airways, constitutively express CD169 (58, 88), an ISG, whose expression can be further induced by type I and type III IFNs (39, 50, 51, 89). While therapeutic use of both type I and III IFNs have been proposed, clinical benefit has proven inconclusive (90–92), presumably related to timing of the IFN response. For instance, a delayed and persistent type I IFN response without resolution has been correlated with disease severity and mortality in patients with COVIID-19 (68). As was recently suggested (93), the pathological outcomes associated with a delayed type I IFN response might be partly due to the type I IFN-induced expression of SARS-CoV-2 entry receptors, such as CD169, in monocytes/macrophages and increased virus uptake and amplification of inflammatory responses.

Treatment of COVID-19 patients with remdesivir has been shown to significantly shorten recovery time and reduce lower respiratory tract infections, despite having minimal impact on viremia (94, 95). Notably, combination therapy with Baricitinib, an anti-inflammatory drug led to further improvement in clinical status and significantly lowered serious adverse events (96). Therefore, targeting macrophage-intrinsic innate immune activation by either blocking macrophage-specific receptors which can mediate SARS-CoV-2 uptake, or through administration of RIG-I/MDA-5 antagonists present a novel therapeutic strategy for combating hyperinflammation in COVID-19 patients. However, further research is needed to better understand the mechanisms of ACE2-independent CD169-mediated SARS-CoV-2 infection of macrophages, the post-entry restrictions to virus replication, and the viral determinants that trigger innate immune activation.

## Materials & Methods

### Ethics statement

This research has been determined to be exempt by the Institutional Review Board of the Boston University Medical Center since it does not meet the definition of human subjects research, since all human samples were collected in an anonymous fashion and no identifiable private information was collected.

### Plasmids

The SARS-CoV-2 S/gp41 expression plasmid was a gift from Dr. Nir Hachoen at the Broad Institute, and has previously been described (74). For ACE2 lentiviral transduction in THP1 cells, we used a lentiviral plasmid expressing ACE2 and the puromycin resistant gene (Addgene #145839, hereafter pLenti-ACE2-IRES-puro). Generation of wildtype and mutant (R116A) human CD169 plasmids (LNC-CD169) was previously described and validated (39). For transduction of primary MDMs, a 3’ LTR-restored lentiviral expression vector (Addgene #101337, hereafter LV-3’LTR) expressing a GFP reporter was used to express ACE2 or CD169. For ACE2 cloning, the NotI-XhoI fragment from pLenti-ACE2- IRES-puro was inserted into the LV-3’LTR backbone. For cloning CD169 into LV-3’LTR vector, a BglII-AgeI fragment from LNC-CD169 was inserted into LV-3’LTR vector. HIV-1 packaging plasmid psPAX2 and VSV-G expression constructs have been previously described (39). All lentiviral vectors (pLKO.1) expressing shRNAs used for knockdown of host proteins were purchased from Sigma.

### Cells

HEK293T cells (ATCC) were maintained in DMEM (Gibco) containing 10% heat-inactivated fetal bovine serum (FBS) (Gibco) and 1% pen/strep (Gibco) (39, 40, 97). Vero E6 cells (ATCC CRL-1586) were maintained in DMEM supplemented with 10% FBS and 100 μg/mL primocin. THP1 cells (ATCC) were maintained in RPMI/1640 (Gibco) containing 10% FBS and 1% pen/strep (50). THP1 cells stably expressing CD169 have previously been described (39). To generate HEK293T/ACE2^+^, THP1/ACE2^+^ and THP1/CD169^+^/ACE2^+^ cells, HEK293T, THP1 or THP1/CD169 cells were transduced with pLenti-ACE2-IRES-puro lentivector and cultured in puromycin-containing media (2 μg/ml). Cells with robust surface expression of ACE2 and CD169/ACE2 (double-positive) were sorted using a MoFlo cell sorter (Beckman Coulter) and cultured in puromycin-supplemented media. All cell lines are routinely tested for mycoplasma contamination and confirmed negative. For THP1 monocyte to macrophage differentiation, THP1 cells were stimulated with 100 nM PMA (Sigma-Aldrich) for 48 hours. Human monocyte-derived macrophages (MDMs) were derived from CD14^+^ peripheral blood monocytes by culturing cells in RPMI/1640 (Gibco) media containing 10% heat-inactivated human AB serum (Sigma-Aldrich) and recombinant human M-CSF (20 ng per ml; Peprotech) for 5-6 days, (55). To generate MDMs expressing ACE2 or overexpressing CD169, cells were co-infected with ACE2 or CD169 expressing lentiviruses (100 ng based on p24^gag^ ELISA per 1x10^6^ cells) and SIV3+ (Vpx expressing) VLPs. ACE2 and CD169 surface expression was determined by flow cytometry 3 days post transduction.

### Viruses

SARS-CoV-2 stocks (isolate USA_WA1/2020, kindly provided by CDC’s Principal Investigator Natalie Thornburg and the World Reference Center for Emerging Viruses and Arboviruses (WRCEVA)) were grown in Vero E6 cells (ATCC CRL-1586) cultured in Dulbecco’s modified Eagle’s medium (DMEM) supplemented with 2% fetal FBS and 100 μg/mL primocin. To remove confounding cytokines and other factors, viral stocks were purified by ultracentrifugation through a 20% sucrose cushion at 80,000xg for 2 hours at 4°C (60). SARS-CoV-2 titer was determined in Vero E6 cells by tissue culture infectious dose 50 (TCID_50_) assay using the Spearman Kärber algorithm. All work with SARS-CoV- 2 was performed in the biosafety level 4 (BSL4) facility of the National Emerging Infectious Diseases Laboratories at Boston University, Boston, MA following approved SOPs.

Generation of SARS-CoV-2 S-pseudotyped lentiviruses expressing spike glycoprotein has previously been described (74). Briefly, HEK293T cells were co-transfected with HIV- 1 reporter plasmid containing a luciferase reporter gene in place of the *nef* ORF and SARS-CoV-2 S (74). To generate ACE2 or CD169 expressing recombinant lentiviruses, LV-3’LTR lentivectors expressing either empty vector, ACE2, or CD169, were co-transfected with psPax2 (HIV Gag-pol packaging plasmid) and H-CMV-G (VSV-G envelope-expressing plasmid) in HEK293T cells by calcium phosphate-mediated transfection (98). Virus-containing supernatants were harvested 2 days post-transfection, cleared of cell debris by centrifugation (300xg, 5 min), passed through 0.45 µm filters, aliquoted and stored at -80 °C until use. Lentivirus titers were determined by a p24^gag^ ELISA (98).

### Infection

For RNA analysis, 1x10^6^ cells (THPI/PMA, MDMs, HEK293T) were seeded in 12-well plates. For smFISH analysis, 1x10^6^ cells were seeded in 6-well plates containing 3-4 coverslips per well. The next day, cells were infected with purified SARS-CoV-2 at the indicated multiplicity of infection (MOI). At indicated time points, the cells were either lysed with TRIzol (for total RNA analysis) or fixed in 10% neutral buffered formalin for at least 6 hours at 4°C and removed from the BSL-4 laboratory for staining and imaging analysis in accordance with approved SOPs. For SARS-CoV-2 S-pseudotyped lentiviral infections of THP1/PMA macrophages or MDMs, 1x10^5^ cells were seeded in 96-well plates, and infected via spinoculation the following day with 10-20 ng (p24^Gag^) of purified S-pseudotyped lentivirus and SIV_mac_ Vpx VLPs (1 hr at RT and 1100 x g), as previously described (55). Incubation with virus was continued for 4 additional hours at 37°C, cells were then washed to remove unbound virus particles, and cultured for 2-3 days. For CD169 blocking experiments, primary MDMs from 3 different donors were pre-incubated with 20 μg/ml anti-CD169 antibody (HSn 7D2, Novus Biologicals) or IgG1k (P3.6.2.8.1, eBioscience) for 30 min at 4°C prior to infection. For anti-spike neutralizing experiments, virus-containing media were pre-incubated with antibodies targeting SARS-CoV-2 spike NTD for 30 min at 37°C prior to infection. At indicated timepoints, cells are lysed, cell lysates were analyzed for luciferase activity using the Bright-Glo luciferase assay kit (Promega), as previously described (36). The SARS-CoV-2 neutralizing antibodies was previously characterized (57) and were a kind gift from Dr. Duane Wesemann at Harvard Medical School.

### S binding

To evaluate SARS-CoV-2 S binding to various THP1 monocytes expressing different surface receptors, approximately 0.25x10^6^ cells from parental THP1 or those expressing wt CD169, mutant CD169 (R116A), ACE2, or both wt CD169 and ACE2 were incubated for 30 min at 4 °C with 2 μg of spike glycoprotein (stabilized) from Wuhan-Hu-1 SARS-CoV-2 containing a C-terminal Histidine Tag, recombinant from HEK293F cells (BEI resources, #NR-52397). This is followed by secondary staining for 30 min at 4°C with APC-conjugated mouse anti-His antibody (BioLegend, #362605, 1:50) or isotype control. Cells were fixed with 4% PFA (Boston Bioproducts) for 30 min, and analyzed with BD LSRII (BD). Data analysis was performed using FlowJo software (FlowJo).

### Immunofluorescence

In brief, the cells were permeabilized with acetone-methanol solution (1:1) for 10 min at - 20°C, incubated in 0.1 M glycine for 10 min at room temperature and subsequently incubated in 5% goat serum (Jackson ImmunoResearch) for 20 minutes at room temperature. After each step, the cells were washed three times in PBS. The cells were incubated overnight at 4°C with a rabbit antibody directed against the SARS-CoV nucleocapsid protein (Rockland; 1:1000 dilution in 5% goat serum), which cross-reacts with the SARS-CoV-2 nucleocapsid protein, as previously described (99). The cells were washed four times in PBS and incubated with goat anti-rabbit antibody conjugated with AlexaFluor594 for 1 hour at room temperature (Invitrogen; 1:200 dilution in blocking reagent). 4’,6-diamidino-2-phenylindole (DAPI; Sigma-Aldrich) was used at 200 ng/ml for nuclei staining. For dsRNA staining (61), anti-dsRNA (Pan-Enterovirus Reagent, clone 9D5, Light Diagnostics, Millipore) antibody was used 1:2 overnight and anti-mouse-AF488 (Invitrogen) 1:200 dilution as secondary antibody with DAPI. Images were acquired using a Nikon Eclipse Ti2 microscope with Photometrics Prime BSI camera and NIS Elements AR software.

### RNA isolation and RT-qPCR

Total RNA was isolated from infected cells using TRIzol reagent (Invitrogen). Reverse transcription (RT) from purified RNAs was performed using oligo(dT)_20_ primer (Superscript III, Invitrogen) or strand-specific RT primers as previously described (100). Target mRNAs were quantified using Maxima SYBR Green (Thermo Scientific), using the primer sets shown in **Table 1**. Primer sequences for GAPDH, IL-6, TNFα, IL-1β, IP-10, IFNλ1, IL-18, Viperin and IFNβ have been described previously (97). The *C_T_* value was normalized to that of GAPDH and represented as a relative value to a ‘mock’ control using the 2^-ΔΔ*C*^ method as described (97, 101).

**Table 1.**
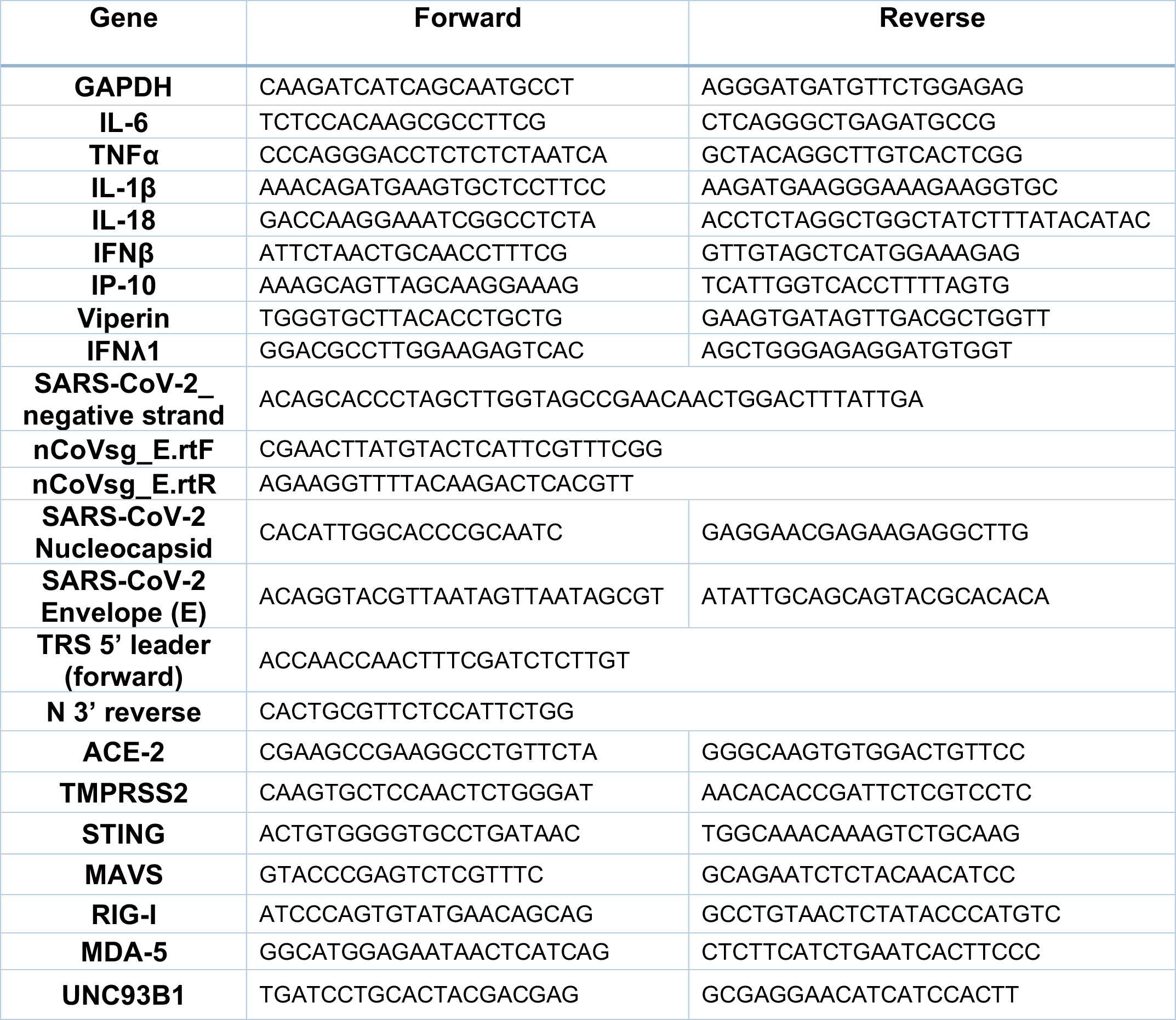
qRT-PCR primers.

### shRNA mediated knockdown

Stable knockdown of host proteins in THP1 cells was carried out by transduction with lentivectors expressing individual shRNAs (pLKO.1, 400 ng p24^Gag^ (as measured by ELISA) per 1 × 10^6^ cells) in the presence of polybrene (Millipore). Cells were washed and cultured for 5-7 days in the presence of puromycin (2 µg/ml, InvivoGen). Selected cells were expanded, and knockdown confirmed and quantified by RT-qPCR or western blotting, prior to any downstream experiments. All shRNA target sequences are listed in **Table 2**.

**Table 2.**
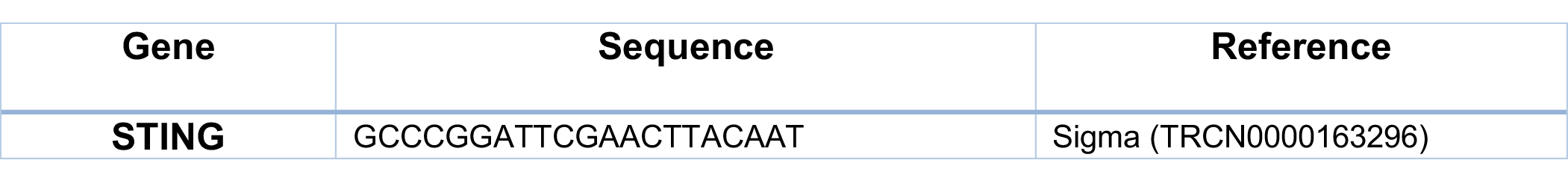
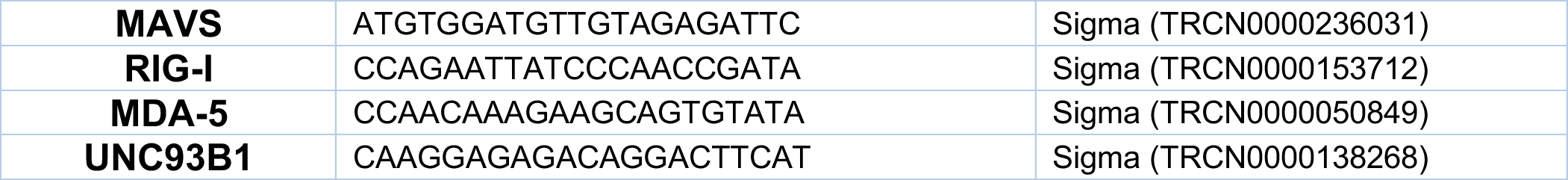
shRNA sequences.

### Flow cytometry

To examine cell surface expression of CD169 or ACE2 in transduced THP1 or primary MDMs, approximately 0.5x10^6^ cells were harvested with CellStripper (Corning), stained with Zombie-NIR (BioLegend, #423105, 1:250) followed by staining for 30 min at 4°C with the following antibodies; Alexa647-conjugated mouse anti-CD169 antibody (BioLegend, #346006, 1:50), Alexa647-conjugated mouse anti-ACE2 antibody (R&D systems, 1:200), or unconjugated goat anti-ACE2 polyclonal antibody (R&D systems, #AF933, 1:200) followed by Alexa488-conjugated chicken anti-goat antibody (Invitrogen, #A-21467, 1:100). Cells were fixed with 4% PFA (Boston Bioproducts) for 30 min, and analyzed with BD LSRII (BD). Data analysis was performed using FlowJo software (FlowJo).

### smFISH

#### Probe designs

smFISH probes used to detect different RNA segments of the SARS-CoV-2 genome (NCBI reference sequence: NC_045512.2) consisted of a set of 48 oligonucleotides, each with length of 20 nt and labeled with different fluorophores (*see Table. 3 in supplementary information*) for target genes and sequences for each of the probe sets). Probes were designed using Stellaris™ Probe Designer by LGC Biosearch Technologies and purchased from Biosearch Technologies. The 3’-end of each probe was modified with an amine group and coupled to either tetramethylrhodamine (TMR; Thermo Fisher Scientific), Texas Red-X (Thermo Fisher Scientific), Quasar 670 (Biosearch Technologies) or Cy5 (Lumiprobe). Coupled probes were ethanol precipitated and purified on an HPLC column to isolate oligonucleotides linked to the fluorophore via their amine groups, as previously described by *Raj et.al.* (62).

#### Hybridization

Cells were cultured on glass coverslips, fixed at appropriate times with 10% neutral buffered formalin and permeabilized with 70% methanol. Coverslips were equilibrated with hybridization wash buffer (10% formamide, 2X SSC), and then immersed in 50 μL of hybridization buffer, which consisted of 10% dextran sulphate (Sigma-Aldrich), 1 mg/mL *Escherichia coli* transfer RNA (Sigma-Aldrich), 2 mM ribonucleoside vanadyl complexes (New England Biolabs, Ipswich, MA), 0.02% ribonuclease-free bovine serum albumin (Thermo Fisher Scientific), 10% formamide, 2X SSC, and conjugated probes with appropriate concentration (25 ng of pooled probes). This hybridization reaction mixture was first added as a droplet onto a stretched-out piece of Parafilm (Bemis in North America, Oshkosh, WI) over a glass plate, and then a coverslip containing the cells was placed faced down onto the droplet, followed by incubation at 37°C overnight in a humid chamber. Following hybridization, the coverslips were washed twice for 10 minutes each in 1 mL of hybridization wash buffer at room temperature. The coverslips were then equilibrated with mounting buffer (2X SCC, 0.4% glucose) and mounted in the mounting buffer supplemented with 1 μL of 3.7 mg/mL glucose oxidase and 1 μL of catalase suspension (both from Sigma-Aldrich) for each 100 μL preparation. After removing the excess mounting medium by gently blotting with a tissue paper, the coverslips were sealed with clear nail polish, and then imaged on the same day.

#### Image acquisition, pre-processing, analysis and mRNA quantification

Images were acquired using Zeiss Axiovert 200M (63x oil immersion objective; numerical aperture 1.4) controlled by Metamorph image acquisition software (Molecular Devices, San Jose, CA). Stacks of images of 16 layers with 0.2 µm interval at 100- to 2,000-milisecond exposure times were used in each fluorescence color channel including DAPI. Two representative coverslips per sample/group were selected and 10-20 regions/fields of interest were imaged. For cell fluorescence intensity measurements, region of interest was drawn manually around each cell using DIC and DAPI channels, then average intensity was measure within the area of each cell using RNA-specific florescence channels.

### Immunoblot Analysis

To assess expression of endogenous or transduced proteins, cell lysates containing 30- 40 µg total protein were separated by SDS-PAGE, transferred to nitrocellulose membranes and the membranes were probed with the following antibodies: mouse anti- TMPRSS2 (Santa Cruz, #515727, 1:1000), mouse anti-Cathepsin-L (Santa Cruz, #32320, 1:1000), goat anti-ACE-2 (R&D systems, #AF933, 1;1000), rabbit anti-STING (Cell Signaling, #13647, 1:1000), rabbit anti-MAVS (Thermo Fisher, #PA5-17256, 1:1000), mouse anti-RIG-I (AdipoGen, #20B-0009, 1:1000), rabbit anti-MDA-5 (Proteintech, #21775-1-AP, 1:1000), rabbit anti-UNC93B1 (Invitrogen, #PA5-83437, 1:1000), rabbit anti-IRF1 (Cell Signaling, #8478S, 1:1000). Specific staining was visualized with secondary antibodies, goat anti-mouse-IgG-DyLight 680 (Thermo Scientific, #35518, 1:20000), goat anti-rabbit-IgG-DyLight 800 (Thermo Scientific, #SA5-35571, 1:20000), or a donkey anti-goat-IgG-IR-Dye 800 (Licor, #926-32214, 1:20000). As loading controls, actin or tubulin expression was probed using a rabbit anti-actin (Sigma-Aldrich, A2066, 1:5000), mouse anti-actin (Invitrogen, #AM4302, 1:5000), or rabbit anti-tubulin (Cell Signaling, #3873, 1;5000). Membranes were scanned with an Odessy scanner (Li-Cor).

### Statistics

All the statistical analysis was performed using GraphPad Prism 9. *P*-values were calculated using one-way ANOVA followed by the Tukey-Kramer post-test (symbols for *p*- values shown with a line) or the Dunnett’s post-test (comparing to mock), symbols for *p*- values shown with a bracket), or a two-tailed paired t-test (comparing two samples, symbols for two-tailed *p*-values shown with a line bracket). Symbols represent, *: *p* < 0.05,

**: *p* < 0.01, ***: *p* < 0.001, ****: *p* < 0.0001. No symbol or ns: not significant (*p* ≥ 0.05).

### Data availability

The authors declare that the data that support the findings of this study are available within the paper and from the corresponding author upon reasonable request.

Table 3 smFISH probe sequences:

See attached excel sheet.

## Acknowledgments

We thank the BUMC Flow Cytometry Core, the Cellular Imaging Core, and Mitchell White, BU for technical assistance. We are grateful to Robert Davey, BU for help with imaging. This work was supported by NIH grants R01AI064099 (SG), R01DA051889 (SG), R01AG060890 (SG), P30AI042853 (SG), R01CA2 27292 (ST), R01AI106036 (YB and ST), R01AI133486 (EM), and R21AI135912 (EM) as well as Fast Grants (EM) and Evergrande MassCPR (EM). The funders had no role in study design, data collection and analysis, decision to publish, or preparation of the manuscript.

## Author Contributions

S.J., J.O., E.M. and S.G. designed the experiments. S.J., J.O, J.B., A.N., E.L., M.L., H.A., Y.B., and S.T., performed the experiments and analyzed the data. S.J., J.O., E.M. and S.G. wrote the manuscript.

## Supplementary Figure Legends

Figure S1. Expression of endogenous TMPRSS2 and Cathepsin-L in human macrophages (**A-B**) Western blot analysis for total TMPRSS2 (**A**) and Cathepsin-L (**B**) expression in wildtype and transduced HT-29 cells (control), HEK293T, THP-1/PMA, and primary MDMs from multiple donors. β-actin was probed as a loading control.

Figure S2. Infection of primary MDMs by S-pseudotyped lentivirus is blocked by anti-SARS-CoV-2 NTD antibodies. SARS-CoV-2 S-pseudotyped lentivirus (20 ng) was pre-incubated with indicated anti- Spike neutralizing antibodies for 30 mins at 37°C, followed by infection of primary MDMs for 3 days. Relative infection quantified by luciferase activity from whole cell lysates. NT: no-treatment with neutralizing antibody. Data are representative of 2 independent experiments, from 3 different donors each. Mock: no virus added, PBS: no-pre-incubation of virus with antibody. The means ± SEM are shown. *P*-values: one-way ANOVA followed by the Dunnett’s post-test comparing to untreated (PBS) control, *: *p* < 0.05, **: *p* < 0.01, ***: *p* < 0.001, ****: *p* < 0.0001.

Figure S3. Expression profiles of human CD169 and ACE2 in THP1 monocytes and primary MDMs. (**A**) Representative flow cytometry profiles of different THP1 cell lines and primary MDMs stained for surface expression of CD169 and ACE2. (**A**) Transduced THP1 cell lines stably expressing wild type (wt) CD169, mutant (R116A) CD169, ACE2, or both wt CD169 and ACE2. (**B**) Untransduced primary MDMs from multiple donors showing differential expression of endogenous CD169. After 5-6 days of macrophage differentiation, cells were either unstained or stained with anti-human CD169 antibody, and surface expression analyzed by flow cytometry. (**C-D**) Representative flow cytometry profiles of primary MDMs transduced with wt CD169 (**C**) or ACE2 (**D**) lentiviruses compared to negative (vector only) control.

Figure S4. Exogenous expression of ACE2 in primary MDMs rescues SARS-CoV-2 replication. Representative immunofluorescence images (20x) of primary MDMs infected with SARS- CoV-2 (MOI=1) and stained for nucleus (DAPI, blue), and SARS-CoV-2 nucleocapsid protein (red), at 24 hpi. Images from primary MDMs overexpressing either CD169 or ACE2 compared to vector-only control were captured and represent at least 3 independent donors. Bar = 25 µm.

Figure S5. CD169 and ACE2-dependent temporal enhancement of SARS-CoV-2 RNAs in THP1/PMA macrophages. Single molecule FISH analysis of viral +gRNA using high fidelity probes in SARS-CoV-2 (MOI=10) infected THP1/PMA macrophages at the indicated timepoints. Representative fields of cells were hybridized at indicated times with 7 sets of smFISH probes labeled with Quasar670 targeting the + strand of SARS-CoV-2 ORF1a (NSP1-3) and N transcripts. Data are representative of 2 independent experiments. Bar = 50µM

Figure S6. Lack of pro-inflammatory cytokine expression in THP1/PMA macrophages infected with SARS-CoV-2 S-pseudotyped lentiviruses. PMA-differentiated THP1 cell lines infected with SARS-CoV-2 S-pseudotyped lentivirus (20 ng) and total RNA harvested at 2 dpi, followed by qRT-PCR analysis. Fold expression of indicated cytokines normalized to mock (uninfected) condition in each group. Data are representative of at least 3 independent experiments. The means ± SEM from 3 independent experiments are shown.

